# Is efficiency of protein incorporations governed by membrane properties? An *α*-synuclein and ceramide-1-phosphate investigation

**DOI:** 10.1101/2025.01.30.635638

**Authors:** Dominik Drabik, Piotr Hinc, Karolina Cierluk, Aleksander Czogalla

## Abstract

In protein-membrane interactions membranes provide an environment that enables proteins to fulfill multiple functions. Yet our knowledge about specificity of protein-membrane interactions is not sufficient. It is our hipothesis that this specificity is governed to a large extend by the properties of membrane itself. This study investigates the protein-membrane interactions in the case of *α*-synuclein (*α*S) and ceramide-1-phosphate (C1P) enriched membranes. The interactions between *α*S and lipids were reported to be governed by various factors, including lipid composition, membrane curvature, charge and fluidity. On the other hand, C1P is anionic lipid and the shape of its molecule is similar to that of PA, which was reported to strongly interact with *α*S. There are three main aspect of this work. First of all interactions between *α*S and C1P enriched membranes is investigated using microscale thermophoresis. Secondly, systematic characterization of membrane systems is performed using both *in vitro* and *in silico* techniques. Eventually, principal component analysis and multiple linear regression are used to determine phenomenological dependency of those interactions in function of membrane properties.

## Introduction

When it comes to protein-protein interactions^1^ or protein-DNA interactions, there is a fair amount of research on this subject. However, some proteins require a lipid membrane to fulfill their function. In summary, looking at the article Scopus database, one can find more than 80 thousand articles regarding protein-protein interactions, more than 14 thousand articles regarding protein-DNA interaction, but only over 2 thousand articles regarding protein-membrane interactions. Furthermore, it was reported that an overview of current applications and developments in high-throughput protein-lipid membrane interactions methods is not yet available^2^. Numerous lipids and proteins are present in the living system, yet we do not have sufficient knowledge about the specificity of their interactions. We now know what the force is that governs that specific protein is abundant at given cellular status while, at the same time, it is lacking in others. To approach this topic, we decided to investigate interactions between *α*-synuclein (*α*S) and various lipid membrane models composed of ceramide-1-phosphate (C1P).

*α*S is a small, intrinsically disordered protein predominantly found in the presynaptic terminals of neurons^3^. It is composed of 140 amino acids with three distinct regions: an N-terminal domain, a central hydrophobic region known as the non-amyloid beta component (NAC), and a C-terminal acidic tail. The versatile structure of *α*S is highly soluble in its native state, but can undergo conformational changes leading to aggregation, which is associated with various neurodegenerative disorders called synucleinopathies, including Parkinson’s disease (PD). It exists in dynamic equilibrium between monomeric, oligomeric, and fibrillar forms, the latter being the main component of Lewy bodies, pathological hallmarks of PD.^4^ It is a multifaceted protein involved in various cellular processes, including synaptic function, protein aggregation, mitochondrial homeostasis, and neuronal survival^5,6^. *α*S has a high affinity for lipid membranes, particularly those enriched in acidic phospholipids such as phosphatidic acid, phosphatidylserine, and phosphatidylglycerol. The interactions between *α*S and lipids were reported to be governed by various factors, including lipid composition, membrane curvature, charge and fluidity^7–9^. Certain lipids, such as cholesterol, have been reported to influence affinity of *α*S and modulate its membrane binding kinetics^10,11^. Interaction with membranes influences its conformation and aggregation propensity, potentially modulating its physiological and pathological functions. One of the possible physiological processes modulated by *α*S – membrane is the regulation of synaptic vesicle dynamics. *α*S plays a crucial role in the trafficking of synaptic vesicles and the release of neurotransmitters.^12^

Its interaction with lipid membranes, particularly synaptic vesicles and the presynaptic membrane, regulates the kinetics of vesicle clustering, docking, and fusion. By modulating membrane curvature and fluidity, *α*S influences the dynamics of the synaptic vesicle, regulating synaptic transmission and neuronal signaling.^13^ Dysregulated interactions between *α*S and lipids have been implicated in the pathogenesis of neurodegenerative diseases, particularly PD and related synucleinopathies. Lipid membranes serve as platforms for *α*S aggregation, facilitating the formation of toxic oligomeric species and fibrillar aggregates characteristic of these disorders.^14,15^ Aberrant interactions between *α*S and lipids can disrupt membrane integrity, leading to cellular dysfunction and neurotoxicity. Oligomeric forms of *α*S may permeabilize lipid bilayers, disrupt ion homeostasis, and induce oxidative stress, ultimately contributing to neuronal degeneration and cell death.^16^

Potentially, one of the lipids that can enhance the interaction of *α*S with membranes, in addition to confirmed phosphatidic acid (PA), is ceramide-1-phosphate (C1P). It is an anionic lipid and the shape of its molecule is similar to that of PA; both lipids have a conical shape as a result of their very small hydrophilic headgroup. C1P consists of a sphingoid base, typically sphingosine, attached to a fatty acid chain through an amide bond. The hydroxyl group of sphingosine is phosphorylated, resulting in the formation of C1P. The fatty acid chain can vary in length and saturation, influencing the physical properties and biological activities of C1P.^17,18^ For a long time, only the known final step in C1P synthesis pathways was through ceramide phosphorylation by ceramide kinase (CERK), an enzyme that was first identified by Bajjalieh et al. and described as ‘calcium stimulated kinase that co-purifies with brain synaptic vesicles’.^19^ However, based on subsequent studies, it was known that there must be a CERK-independent C1P synthesis pathway.^20^ In recent years, it has also been discovered that C1P can be produced by ceramide phosphorylation by the enzyme diacylglycerol kinase zeta (DGK*ζ*), which until now was considered an enzyme that exclusively phosphorylates diacylglycerols, resulting in the production of phosphatidic acid.^21^ C1P acts as a signaling molecule involved in various cellular processes, including cell proliferation, survival, migration, and inflammation. It can exert its effects through interactions with specific protein targets or by modulating the properties of cellular membranes by influencing membrane curvature and fluidity, which may affect membrane trafficking, vesicle formation, and organelle morphology.^17,18,22^

In this article, we have investigated the interaction of *α*S with lipid membranes enriched with two distinct subpopulations of ceramide-1-phosphate in the acyl chain structure (namely C18:1 C1P and C16:0 C1P) in the presence and absence of cholesterol in the membrane. This was performed by microscale thermophoresis (MST). Both the Hill coefficient and the effective concentration parameters (such as *EC*_50_) were determined. This was followed by a biophysical characterization of the membrane containing C1P. We have combined in vitro and in silico methodologies such as Langmuir monolayers, flicker-noise spectroscopy, and molecular dynamics simulations to characterize membranes enriched with C1P and cholesterol. As a result, we have a dataset of parameters including experimentally obtained area per lipid (APL), bending rigidity, compressibility modulus, and free Gibbs energy of mixing, as well as computationally obtained membrane thickness, diffusion coefficient, interdigitation, tilt energy, and defect analysis. Finally, we implemented both a principal component analysis (PCA) and a structure-activity relationship (SAR)-like approach to establish which of those membrane parameters have correlated with protein-membrane binding parameters. Taking the Hill coefficient as an indicator of interaction cooperativity and *EC*_50_ as an indicator of protein-membrane affinity, we were able to designate candidates that most likely are responsible for those aspects. As such approaches provide more reliable results for a more numerous sample, we added to our analysis membranes with two most structurally similar subpopulations of phosphatidic acid (16:0-18:1 POPA and 16:0-16:0 DPPA). By juxtaposing the C1P data with the previously obtained membrane characteristics of the PA-enriched membrane^23^, we were able to obtain more general conclusions regarding possible relationships between protein-membrane interactions and membrane properties.

## Results & Discussion

### Interaction of αS with membranes containing C1P

For membranes composed of phosphatidylcholine (POPC) with the addition of 30 mol% cholesterol, the binding was monophasic (Figure 1A), with an *EC*_50_ of 348.9 ± 19.5 (Figure 1B) µM and a Hill coefficient of (2.79 ± 0.44) (Figure 1C)). Membranes made solely of POPC without cholesterol also exhibited monophasic binding but with a slightly higher, but not statistically significant (comparing to POPC only system), *EC*_50_ of 392.82 ± 43.27 µM and a similar Hill coefficient of (2.80 ± 0.79).

**Figure 1.**
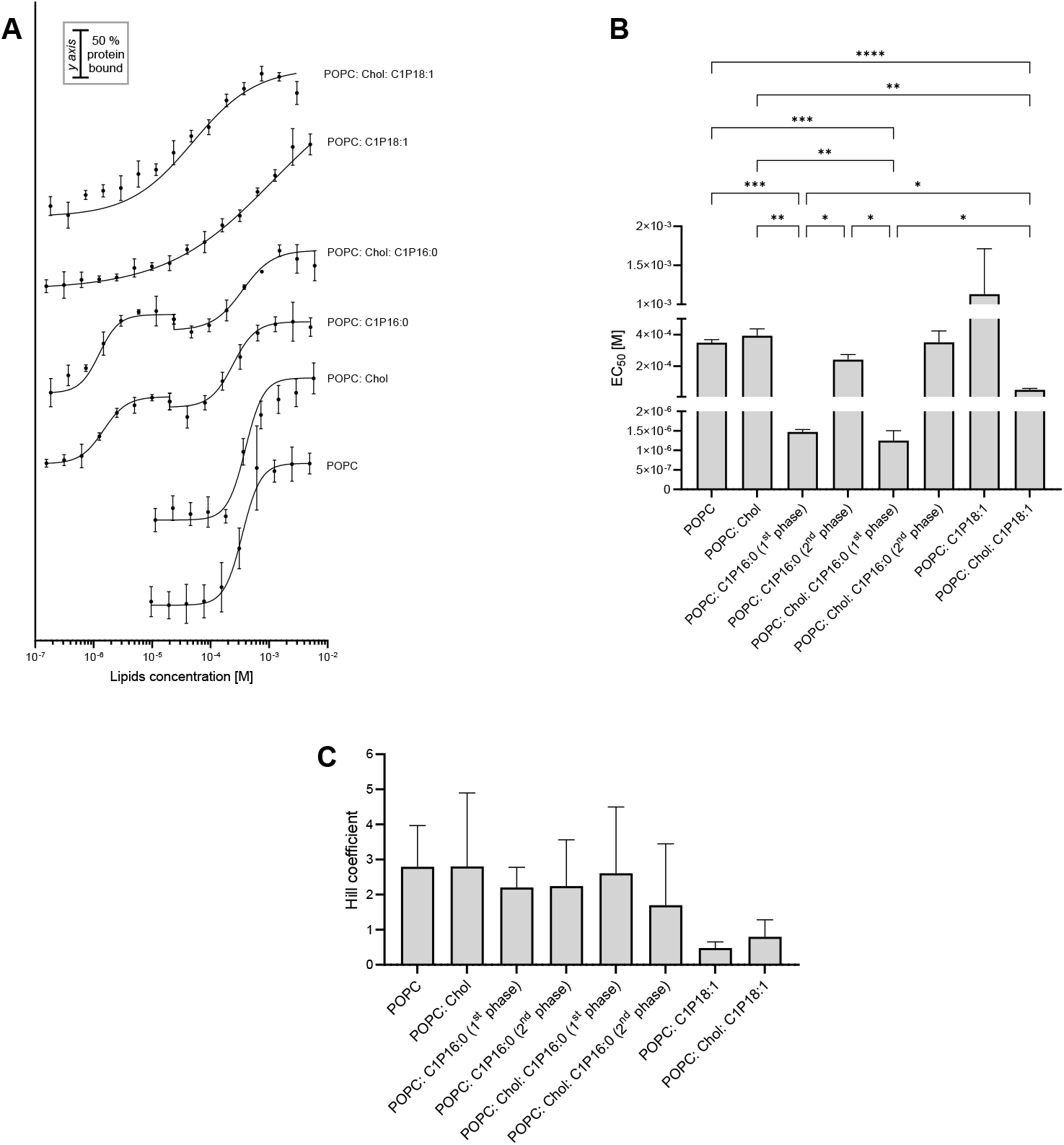
Analysis of interaction between *α*-synuclein and membranes containing different sub-population of C1P in the presence and absence of cholesterol. (A) Binding curves of protein-membrane interactions. (B) Values of *EC*_50_ calculated from fitting of binding curves. Comparisons between samples that were not indicated were not statistically significant. (C) Values of Hill coefficient calculated from fitting of binding curves. There were no statistically significant differences between analyzed groups. * p<0,05; ** p<0,01; *** p< 0,001; **** p<0,0001.

For membranes containing 20 mol% saturated C1P (16:0), the binding exhibited a biphasic nature, indicating the presence of two distinct binding sites or modes. The first binding phase showed an affinity with an *EC*_50_ of 1.47 ± 0.06 µM and a Hill coefficient of 2.2 ± 0.3, suggesting positive cooperativity. The second phase showed a much lower affinity with an *EC*_50_ of 241.2 ± 32.6 µM compared to the first phase with a Hill coefficient of 2.24 ± 0.54, also indicative of positive cooperativity. The presence of cholesterol in membranes containing 20 mol% of saturated C1P (16:0) slightly altered the binding profile. The biphasic nature persisted, but *EC*_50_ for the second phase increased to 350.7 ± 72.4 µM, with a Hill coefficient of 1.70 ± 0.71. This could suggest a decrease in binding affinity and cooperativity compared to the cholesterol-free system; however, these changes are not statistically significant. The first phase maintained an affinity with an *EC*_50_ of 1.25 ± 0.25 µM and a Hill coefficient of 2.61 ± 0.85, reflecting a strong cooperative interaction, and also did not show statistically significant differences compared to the first phase of binding to the cholesterol-free system. For membranes containing 20 mol% unsaturated C1P (18:1), a monophasic binding curve was observed, with an *EC*_50_ of 1130 ± 583 µM and a Hill coefficient of 0.48 ± 0.05. The Hill coefficient value below 1 indicates negative cooperativity and much lower binding affinity compared to all the systems described above - even control systems without C1P. The addition of cholesterol to membranes containing 20 mol% unsaturated C1P (18:1) resulted in a monophasic binding curve with a significantly increased (p<0.01 - compared to cholesterol-free system) binding affinity with *EC*_50_ of 49.4 ± 9.6 µM and a Hill coefficient of 0.80 ± 0.14, suggesting a slight or no increase in cooperativity compared to the unsaturated C1P alone, as this change shows no significant difference.

The results highlight that the interaction of *α*-synuclein with lipid membranes is significantly influenced by the lipid composition. Specifically, the saturation level of the C1P acyl chain and the presence of cholesterol play crucial role in modulating binding affinity and cooperativity. The observed biphasic nature in some interactions suggests the existence of multiple binding sites or binding modes, possibly due to domain formation in the membrane, reflecting the complexity of the *α*-synuclein-lipid interaction landscape.^24^ The results indicate that synuclein exhibits the highest affinity for membranes containing the saturated C1P sub-population, with this binding showing high cooperativity, while cholesterol has no impact on the interaction parameters (*EC*_50_ and Hill coefficient) in this system. In the case of the unsaturated C1P sub-population, we observe both a reduction in protein affinity for the membrane and an increase in this affinity (compared to control systems without C1P), which is regulated by the presence of cholesterol in the membrane. The presence of cholesterol in membranes containing C18:1 C1P enhances the protein’s affinity for the membrane. Interactions with membranes containing C1P (18:1) show negative cooperativity or noncooperative binding events. This indicates that the presence of C1P/cholesterol in the membrane generates certain physicochemical properties that modulate the protein’s affinity for the membrane, the cooperativity of this binding, and the number of binding sites within the protein or membrane. To this end, we followed with the characterization of the physicochemical properties of the investigated membrane systems.

### Physicochemical characterization of C1P membrane systems

All lipid membrane systems with C1P were subjected to detailed characterization of their structural and mechanical properties using both experimental and numerical techniques. Langmuir monolayer technique was employed to determine the area per lipid (APL), excess free energy of mixing (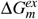), and monolayer compressibility (*C*_*s*_), and the results were supplemented with similar parameters obtained from MD simulations. Obtained values of determined APL from both experimental Langmuir monolayer and MD simulations are presented in Figure 2A (numerical values are presented in Table S1 ). For the monolayer system composed only of C1P (16:0), the obtained value of APL was equal to (61 ± 2) Å^2^ (taken at pressure 33 mN/m, at T=(22.0 ± 0.5) °C, compression ratio 10 mm/min which corresponds to compression rate of 9.7 Å per molecule per minute, pH 7.4). Interestingly, this value is significantly higher than that reported in other studies. It was reported to be equal to 46 Å^217^, but with a slightly lower temperature T = 20 ^*°*^C and a lower compression rate equal to 1.8 Å^2^ per molecule per minute. In another study, it was reported between 39 Å^2^ and 41 Å^225^, for pH 9 and 4, respectively, lower temperature T=20 ^*°*^C and with compression rate equal to 3 Å per molecule per minute. It is our understanding that the APL of C1P (16:0) strongly depends on the compression rate and can lead to such discrepancies between different conditions. To this end, we put an additional focus on maintaining the same compression rates between all of the investigated systems. For a monolayer system composed only of C1P (18:1), the APL was equal to (42.6±1.0) Å^2^. The literature data on C1P (18:1) is scarce, the best comparison could be made with D-erythro-C18 Ceramide, for which APL was reported to equal 39 Å^2^ .^26^ A comparison between obtained and theoretically calculated values from corresponding mono-lipid systems is presented in Supporting information section 1.1 . Interestingly, we also observed the compression effect of monolayers as shown by the negative values of 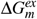 in the case of the POPC:C1P (18:1) system in Figure 2B (see Supporting information section 1.2 for more information ).

**Figure 2.**
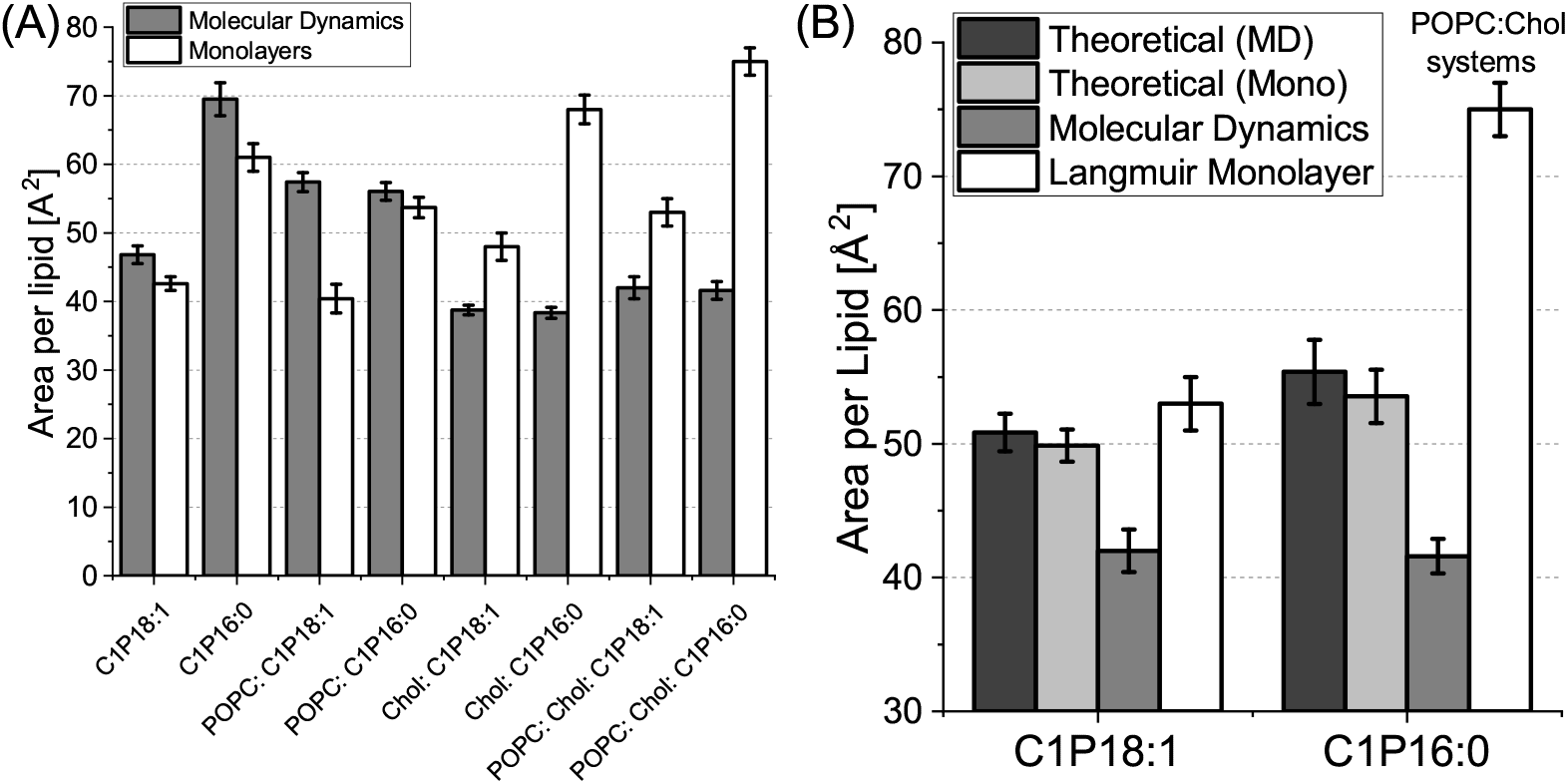
(A) Comparison between APL (area per lipid) determined from molecular dynamics simulations and Langmuir monolayers measurement. (B) Calculated value of excess free energy of mixing for investigated systems.

Finally, the monolayer compressibility is presented in Figure 3A for all investigated systems. Area compressibility, for membrane systems, was also determined using MD (see Table S2) . A general trend can be seen that membranes with C1P (16:0) have higher compressibility than their counterparts with C1P (18:1). Only in the case of membrane systems with POPC the value for POPC:C1P (18:1) was higher than POPC:C1P (16:0). This is true for both monolayer compressibility *C*_*s*_ and membrane stretching compressibility *K*_*A*_. Interestingly, the values of *K*_*A*_ for membrane systems are much higher than the simple twice value *C*_*s*_ of a monolayer system. This indicates that the organization of molecules in membrane systems is significantly different from that of monolayer ones in this case. Nevertheless, in each of the investigated systems, the addition of cholesterol increases these parameters. This increase is much smaller in monolayer systems (around ∼ 1.5 times) but significantly higher in membrane systems (even up to ∼ 15 times). The second parameter describing the mechanical behavior of the lipid membrane is the bending rigidity coefficient *κ* - a parameter that describes the effective energy cost of bending a membrane. It was determined using both experimental Flicker-noise spectroscopy and MD simulations. The results are presented in 3B. There, bending rigidity exhibits a different trend than compressibility - the value is inversely higher in systems with C1P (18;1). In addition, cholesterol dramatically increases the energy that is required to bend the membrane. A similar effect was observed in the case of PA lipid-containing systems and was explained with a strong synergistic effect of both PA and Chol on membrane mechanics, which does not occur in the case of two-component membranes^23^. The parameters established for individual vesicles are presented in Figure S4 . Tilt energy was also established using MD simulations. This empirical parameter can be understood as the measure of forces associated with the tilting of a lipid in the membrane – the more restricted the possibility of tilt movement of the lipid, the higher value of this parameter (see 3C). One can see significant tilt energy in the case of C1P (16:0) homogeneous system, which could indicate tilting effect - exhibition of strong self-limiting movement tendency - associated with gel phase. This could explain the extreme behavior for this system in the case of all the parameters determined. Readers interested in biophysical basis of lipid membrane mechanical properties could follow several good reviews in this subject^27,28^.

**Figure 3.**
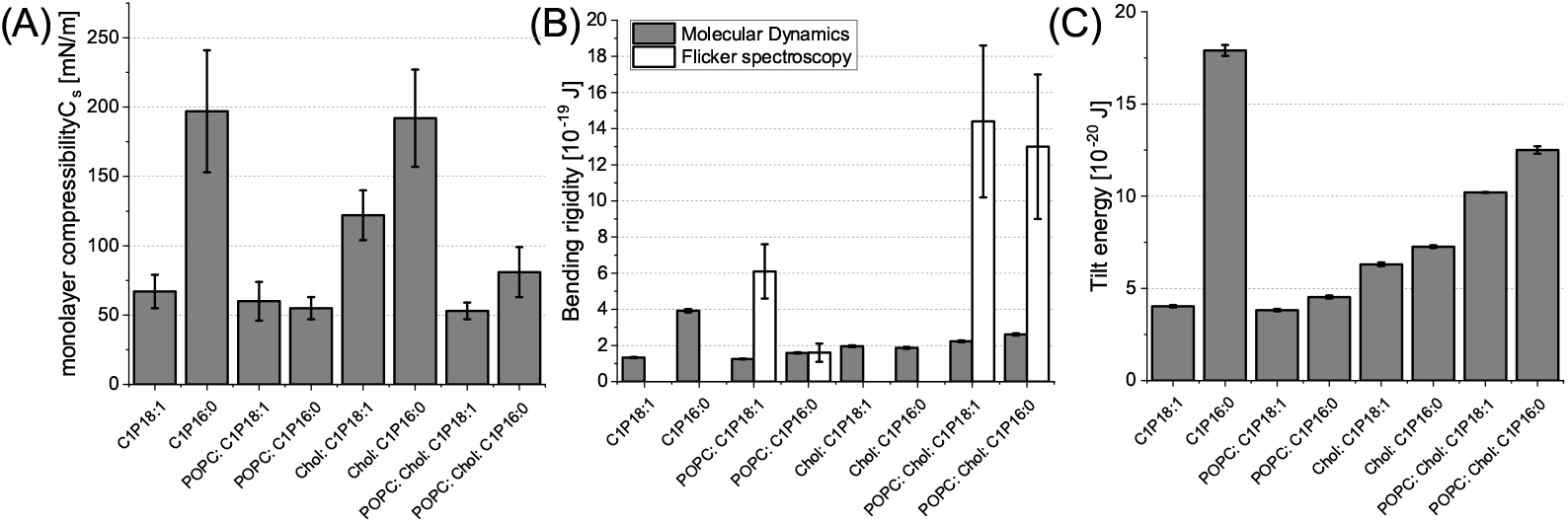
Mechanical parameters obtained from Langmuir monolayers and molecular dynamics simulations of membrane system. Values of (A) monolayer compressibility, (B) bending rigidity and (C) tilt energy are presented for all investigated systems.

MD simulation can give access to parameters that can be quite challenging to establish using experimental techniques. Those parameters are membrane thickness, lateral diffusion of lipids, order parameter and interdigitation. The last one can be understood as the overlap of acyl chain regions between two leaflets and gives information about interleaflet interactions or even possible coupling. In addition, a detailed analysis of defects on the membrane surface can be performed. The numerical values of the parameters obtained, as well as the information on statistical significance, details of individual simulations, and detailed graphs of individual experiments are presented in Table S2, Figure S5.A . An interesting fact is that while a homogeneous membrane composed of C1P (16:0) has a strong effect by itself on all parameters, when C1P (16:0) is present in a heterogeneous membrane, the effect can be opposite. For example, the MT of the homogeneous C1P (16:0) membrane is lower than the C1P (18: 1), but when comparing POPC:C1P (16:0) and POPC:C1P (18:1), the latter is lower. A similar trend is observed in the mixtures C1P:cholesterol and POPC:cholesterol:C1P. This behavior is visible in the case of APL, mechanical parameters, and also defects (Figure 4A). Only in the case of lateral diffusion, this tendency is broken - membranes with C1P (16:0) have, in all investigated mixtures and homogeneous setups, lower diffusion than their counterparts enriched with C1P (18:1).

**Figure 4.**
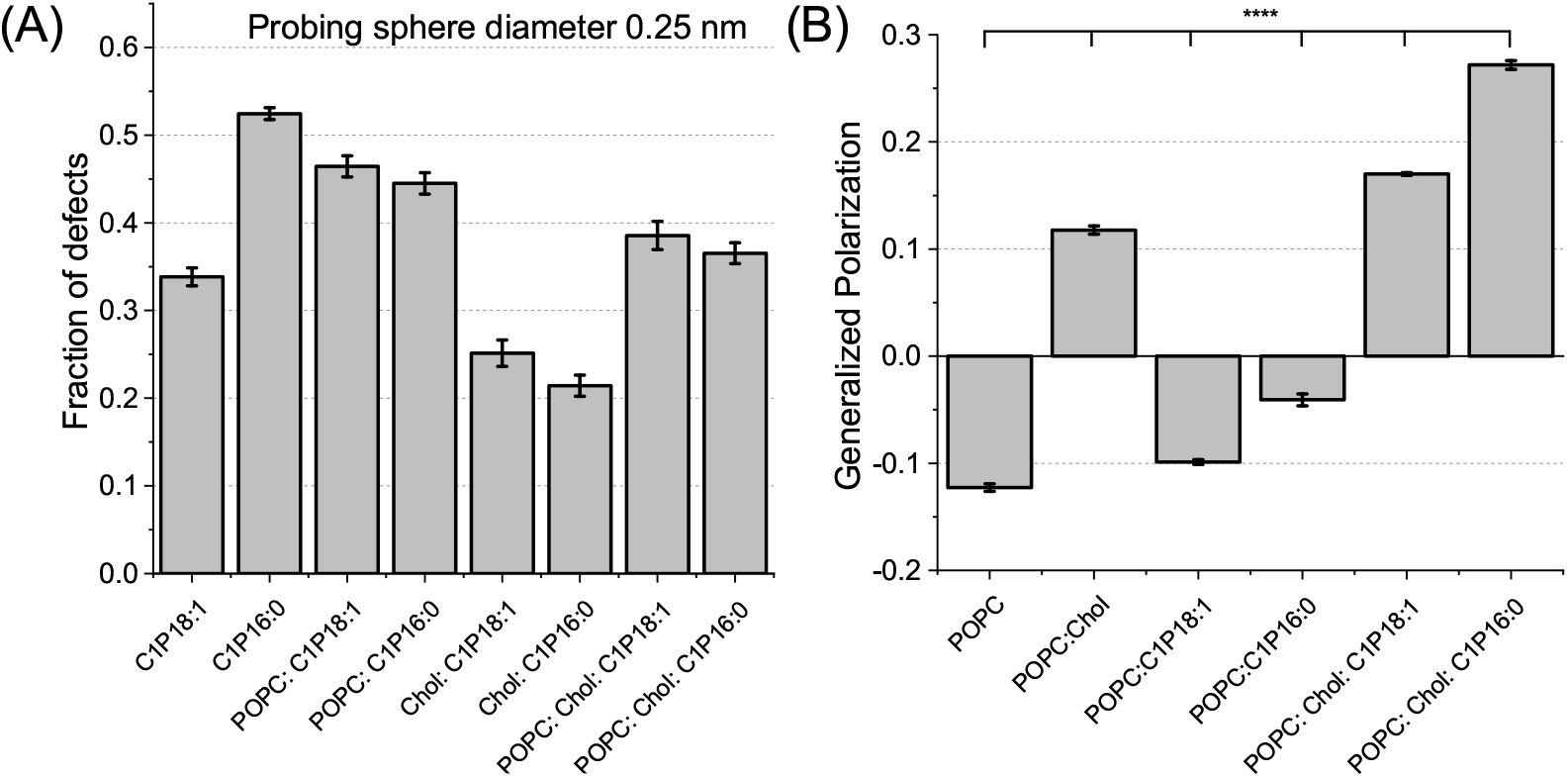
(A) Calculated fraction of defects in membrane system for spherical probe of diameter 0.25 nm. (B) Calculated values of GP. **** p<0,0001.

Addition of C1P to membranes, both in the presence and absence of cholesterol, results in an increase in generalized polarization (GP) values (Figure 4B), which reflects an increase in lipid order within the membrane. The addition of the saturated form of C1P (16:0) to cholesterol-free membranes leads to a (67±1)% increase in GP (p<0.001) compared to the control system without C1P, and a (131±10)% increase (p<0.001) when C1P is added to cholesterol-containing membranes, compared to the cholesterol-containing control system without C1P. In contrast, the addition of the unsaturated form of C1P (18:1) to cholesterol-free membranes results in a (19±2)% increase in GP (p<0.001) compared to the control system without C1P, and a (44±5)% increase (p<0.001) when C1P is added to cholesterol-containing membranes, compared to the cholesterol-containing control system without C1P. Averaged fluorescence emission spectra are presented in Figure S5.B . Based on these results, it can be concluded that C1P (16:0) induces a greater increase in GP compared to C1P (18:1), which is consistent with the fact that saturated acyl chains are introduced into the membrane. Interestingly, in both C1P sub-populations, the increase in GP caused by C1P incorporation into the membrane is approximately twice as high in membranes containing cholesterol. Unlike ceramide, which tends to alter bilayer integrity and can induce hexagonal phases, C1P is more likely to favor lamellar structures similar to sphingomyelin^29,30^. This structural preference suggests that C1P may stabilize membrane organization in the presence of cholesterol. Cholesterol is known to interact with sphingolipids, including C1P, through hydrogen bonding and van der Waals forces, which contribute to the formation of liquid-ordered domains within the membrane^29,31^. These interactions can enhance the acyl chain order of the surrounding lipids.^32^

### Modeling of binding descriptive parameters with membrane properties

Finally, we implemented a structure-activity relationship -like approach based on multiple line regression (MLR) to establish which of the determined membrane parameters correlates with protein-membrane binding parameters. Specifically, we hypothesize that the incorporation of the investigated protein is governed to large extent by the membrane properties of the given system. To this end, a principal component analysis was performed to investigate how membrane parameters are grouped and what is their relation to binding protein parameters such as Hill coefficient and half maximum effective concentration *EC*_50_. However, this type of study allows for more general conclusions when a higher number of statistical samples are presented. To this end, as we mentioned in Introduction section, we added to our analysis membranes with two most structurally similar sub-populations of phosphatidic acid (16:0-18:1 and 16:0-16:0 PA)^23^. The additional results of protein binding to PA-enriched along the numerical values of C1P-enriched membranes are presented in Table S3 . The *α*S protein was previously reported to have a strong affinity towards such membranes^33,34^, which justifies the selection. These results are presented in Figure 5. One can see that the parameters are grouped into three main subsets. One set of parameters (yellow) is related to the probed defects and the thickness of the membrane together with *EC*_80_. The second (green) consists of mechanical parameters such as bending rigidity, *C*_*s*_, mixing constant, *K*_*A*_, order parameter and experimentally determined APL. Finally, the third subset (red) consists of loosely connected simulation parameters such as computationally determined lateral diffusion, interdigitation, polar ratio and APL. Interestingly, most protein-related parameters are not specifically located in any of those subsets. In the case of the Hill coefficient that is localized in the center, this indicates that only a combination of parameters from some of the groups could model it reasonably well. In the case of *EC*_50_, as well as *EC*_20_, it seems to be mostly associated with the yellow set of parameters (defect one) with a small influence of the red one.

**Figure 5.**
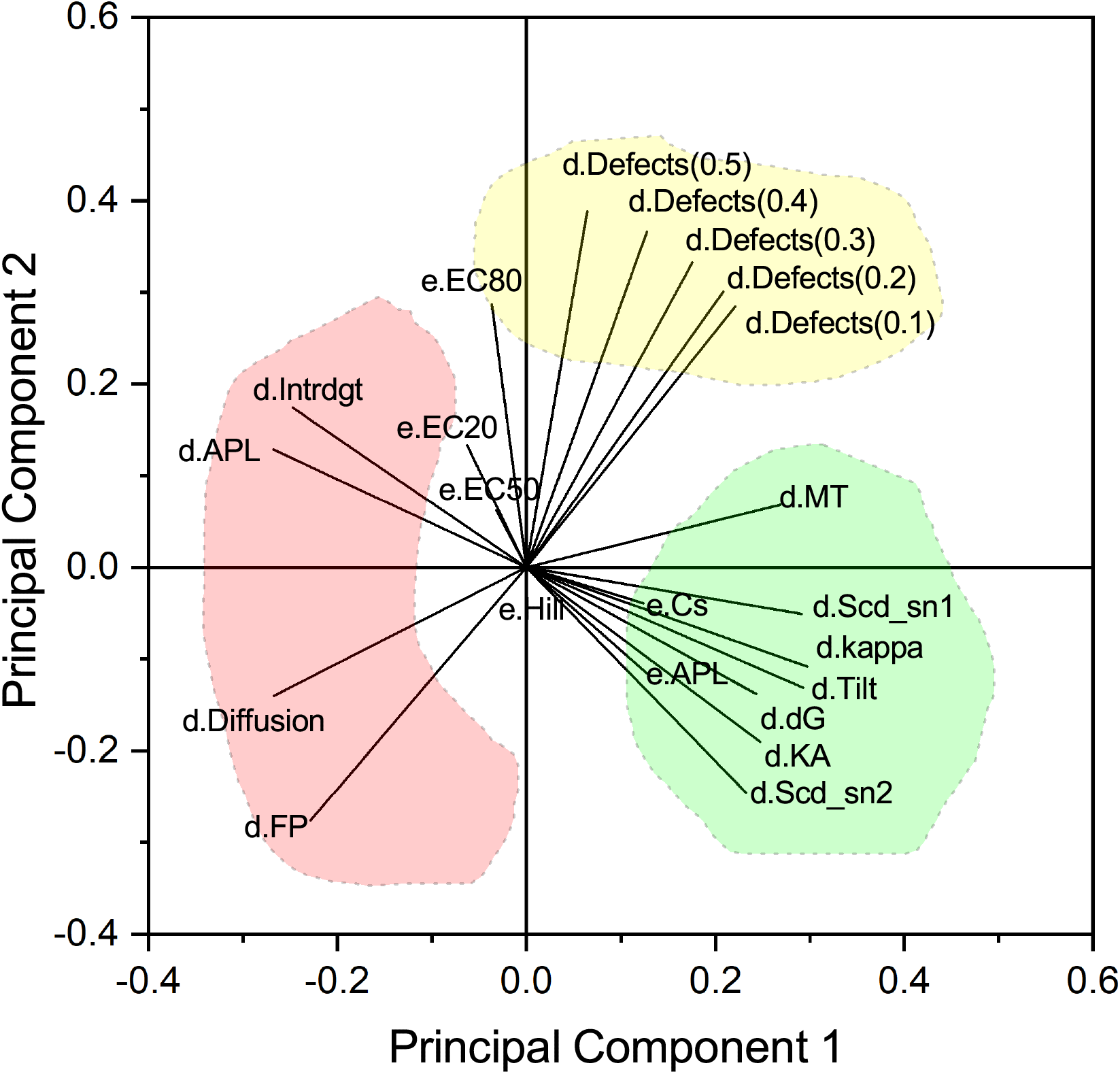
Principal component analysis of the whole dataset. To increase clarity of variables used, the variable is either prefixed with e or d indicating value from experiments or molecular dynamics. The detailed explanation of abbreviations is presented in Table S4 for clarity.

To model both protein parameters with membrane properties, we adapted a multiple linear regression model. This allowed for the determination of the most accurate membrane properties that can then be used to obtain experimentally determined protein parameters. As only a limited number of various membranes were investigated in this study, we decided to consider no more than 3 membrane properties in this study. To further increase the precision and general character of our analysis, we also characterized two selected systems with 10% C1P, specifically: POPC:C1P (16:0) and POPC:Chol:C1P (18:1). Each of the membrane parameters determined was additionally subjected to various mathematical operations to include possible correlation/dependency. A final number of descriptors was equal to 81, which are listed in detail in the Table S4 . A number of training sets (characterized membrane systems) was equal to 10 while the number of verification sets was equal to 2. As a result, we investigated 85320 models from which we selected the most accurate based on the sum of residues of both training and verification sets. We started by modeling the Hill coefficient. As mentioned previously, the Hill coefficient can be interpreted as an indicator of cooperative binding. It is often used to describe the binding of ligands to macromolecules. In general, the binding of a ligand to a macromolecule is enhanced when other ligands are present on the same macromolecule. However, as our design of the experiments differs from classical, it should be noted that the Hill coefficient can not be strictly an indicator of only cooperativity, but also liposome-protein binding phenomena. In Table 1 three best correlated models for Hill coefficient are presented. The juxtaposition of experimental and modeled values of the best model, as well as assessment model efficiency on training set are presented in Figure 6. The Phenomenological equation obtained from the best model #1 was *Hill*_*Coe f f*_ = −1.088 * *log*(*kappa*_*e*_*xp*) − 0.531 * *log*(|*Mxc*|) + 9.103 * *sqrt*(*Sn*1) + 0.799. Similar graphs for two other models are presented in Supporting Information Figures S10 .

**Table 1.**
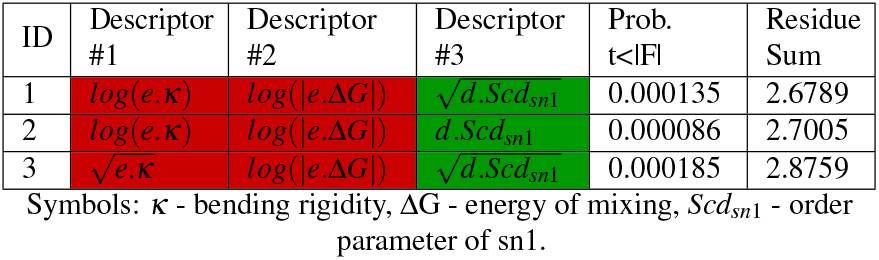
List of three models for Hill Coefficient determination with lowest residue for both training and validation along descriptors of those models and their probability. Green background reflects positive correlation while red - negative correlation.

| ID | Descriptor #1 | Descriptor #2 | Descriptor #3 | Prob. $t < \vert F\vert $ | Residue Sum |
| --- | --- | --- | --- | --- | --- |
| 1 | $\log(e.\kappa)$ | $\log(\vert e.\Delta G\vert )$ | $\sqrt{d}.Scd_{sn1}$ | 0.000135 | 2.6789 |
| 2 | $\log(e.\kappa)$ | $\log(\vert e.\Delta G\vert )$ | $d.Sc d_{sn1}$ | 0.000086 | 2.7005 |
| 3 | $\sqrt{e.\kappa}$ | $\log(\vert e.\Delta G\vert )$ | $\sqrt{d}.Sc d_{sn1}$ | 0.000185 | 2.8759 |

Symbols: $\kappa$ - bending rigidity, $\Delta G$ - energy of mixing, $Scd_{sn1}$ - order parameter of sn1.

**Figure 6.**
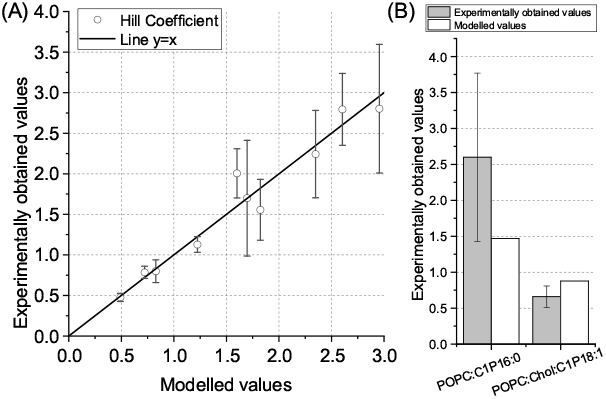
(A) The juxtaposition of experimentally determined values of Hill Coefficient (OY axis) and modeled values (OX axis). The black line represents perfect agreement between two datasets. (B) Comparison between experimentally obtained values of Hill coefficient (gray) and modeled values (white) in the training set.

Interestingly, each of the three best models depended on the same membrane properties with a small difference in the presence of the root of the selected parameters. Specifically, we observed a strong negative dependence (-1.088 in model # 1) on experimentally determined bending coefficients. This was followed by an even stronger dependence on averaged ordered parameters of the *Sn*_1_ lipid tail. Finally, the third parameter was the logarithmic value of the experimentally determined excess free energy of mixing. This indicates that cooperativity strongly depends on the ordering of lipid molecules in the membrane: the more disordered the lipid tail, the easier it is for the protein to penetrate if at least some proteins are already in the system. Indeed, the order of the lipid tails in membranes can impact the binding cooperativity of intrinsically disordered proteins (IDPs) by influencing the accessibility of binding sites on the membrane surface. Studies have shown that the lipid tails order can modulate the conformational dynamics of IDPs upon membrane binding, affecting the cooperativity of their interactions with membranes^35^. Furthermore, we can observe a negative dependence on the rigidity of the bending, so the higher the cost of the bending of the membrane, the lower the cooperativity of the *α*S protein binding to the membrane. Indeed, membrane bending can play a crucial role in IDP-membrane interactions. Change of membrane bending, induced by proteins like epsin that insert helices into lipid bilayers, can support structural transitions in membranes, such as the flat-to-dome transition, and stabilize membrane curvature, impacting the binding of IDPs to membranes^36,37^. The bending of membranes can create spatial constraints that influence the binding modes and cooperativity of IDPs with the membrane possibly through the generation/modification of binding sites, as these defects are enhanced at positive membrane curvature due to the mismatch between the membrane curvature and the intrinsic curvature of lipids^38,39^. Interestingly though, we did observe the negative dependency of bending on the cooperativity of *α*S protein, which implies that stabilization of curvature is more crucial than generation of binding sites for the cooperativity. Finally, we also observed a negative dependence on the excess free energy of mixing. In general, this property is a measure of energy change due to mixing – to consider a process spontaneous, the energy of mixing should be negative, otherwise, an external energy is required for the mixing of membranes. In the case of an ideal miscible system, it should equal zero. As a result, we can treat this parameter as an indicator of possible heterogeneity distribution of lipids in the membrane. This suggests that the higher the cost of the membrane system to mix, the phenomenon of positive cooperativity during protein binding to the membrane decreases.

A similar analysis was performed for the parameters *EC*_50_. However, since *EC*_50_ are not linearly distributed values, we were unable to obtain the modeling of *EC*_50_ because we found an issue of lack of data uniformity. To overcome this issue, we modeled *log*10(*EC*_50_), as it provided us with more linear distributions of the experimental data. This allowed for significantly better validation and *R*^2^ parameters. The value of *EC*_50_ in our case is simply understood as the concentration of lipids that produces half of the maximal response in MST experiment (lipids concentration at which half of protein molecules is bound). The lower the *EC*_50_ value was observed, the lower the concentration of lipids was required to produce 50% of the maximum response, suggesting higher affinity. To this end, we will use the *EC*_50_ value as an indicator of *α*S potency to bind to a given membrane system. In Table 2 the three best correlated models are presented. The juxtaposition of experimental and modeled values of the best model, as well as the efficiency of the assessment model in the training set, is presented in Figure 7. The phenomenological equation obtained from the best model #1 was *EC*_50_ = 0.265 * *sign*(*e*.Δ*G*) − 35.08 * *sqrt*(*d*.*Scd*_*sn*1_) − 14.297 * *e*^*e*.*GP*^ − 2.96. Similar graphs for two other models are presented in Supporting Information Figures S11 . However, it should be noted that while the best model did well in juxtaposition of experimental vs. modeled values, the estimation of the training set did, at least in the case of POPC:Chol:C1P (18:1) unsatisfactory. However, of the three models discussed, only #3 estimated the training set correctly, though at the expense of juxtaposition.

**Table 2.** List of three models for Hill Coefficient determination with lowest residue for both training and validation along descriptors of those models and their probability. Green background reflects positive correlation while red - negative correlation.

| ID | Descriptor #1 | Descriptor #2 | Descriptor #3 | Prob. $t < \vert F\vert $ | Residue Sum |
| --- | --- | --- | --- | --- | --- |
| 1 | $sign(e.\Delta G)$ | $\sqrt{d}.Scd_{sn1}$ | $e^{e.GP}$ | 0.00028 | 2.2364 |
| 2 | $d.Def_{0.3}$ | $\log(d.Def_{0.5})$ | $(d.Intrg)^3$ | 0.00210 | 2.5473 |
| 3 | $\sqrt[3]{d}.Def_{0.4}$ | $\log(d.Def_{0.5})$ | $\log(d.MT)$ | 0.00460 | 2.3486 |

Symbols: $\Delta G$ - energy of mixing, $Scd_{sn1}$ - order parameter of sn1, $GP$ - generalized polarization, $Def$ - defects, $Intrg$ - interdigitation, $MT$ - membrane thickness.

**Figure 7.**
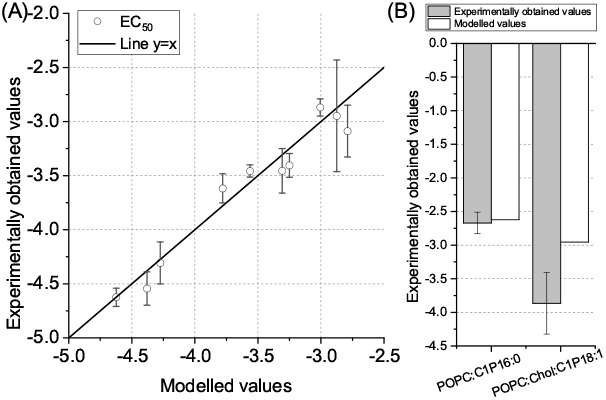
(A) The juxtaposition of experimentally determined values of *EC*_50_ (OY axis) and modeled values (OX axis). The black line represents perfect agreement between two datasets. (B) Comparison between experimentally obtained values of *EC*_50_ (gray) and modeled values (white) in the training set.

Unlike the Hill Coefficient, *EC*_50_ modeling depends mostly on properties obtained from simulations of lipid membranes with small exception for GP and sign of mixing constant values in model #1. Furthermore, while the residual sum was of similar value to the previous case of the Hill coefficient, the dependency of modeled vs experimental values was more deviated from linear character. There were a few outliers in juxtaposition for models #2 and #3. Only in the case of model #1 the outliers were not present, but the estimation for one of the membranes in the training set was significantly worse. Nevertheless, also in this case all of the best models were mostly based on the same properties of membranes. We observe a strong correlation in the case of model #1 with GP. This suggests that domain formation does influence the incorporation of investigated protein. Furthermore, we observe a strong correlation with a fraction of defects observed with spherical probes of radii 0.3-0.5 nm in models #2 and #3. It was particularly strong in the case of model #3 ( ∼ 42), but also not lacking in the second one ( ∼ 24). This was followed by a second parameter - a logarithm of the fraction of defects observed with spherical probes of radius 0.5 nm, though interestingly in each of the models, this was a negative correlation (from -4 to -7). This indicates that the potency of incorporation of *α*S into membranes is mostly governed by the size of defects. Simply put, protein incorporates when the acyl-chain region is sufficiently exposed (0.3 - 0.4 nm) to allow the protein for incorporation, but not to exposed (0.5 nm). This is indeed supported by literature. The size of defects in membranes plays a crucial role in influencing the binding of *α*S and other IDPs. These defects, which are locations on the membrane surface where the hydrophobic interior is exposed to the solvent to accommodate high curvature, are particularly relevant for proteins containing amphipathic helices like *α*S^40^. Studies have shown that *α*S preferentially binds to highly curved, negatively charged membranes by identifying lipid packing defects^41^. Interestingly we did not observe any of the investigated parameters describing curvature of the membrane in our models. The reason for this might be, that it is indeed defects that are crucial for the membrane rather than the curvature itself, which is just a means for higher defects presence. The interaction between IDPs and membranes is intricate, with the amphipathic helix of *α*S and other proteins detecting and binding to these defects to facilitate membrane association. The defect size in membranes can significantly impact the binding affinity and dynamics of IDPs like *α*S. Larger defects on curved membranes have been found to promote the folding of binding domains, with the defect structure responding to the bound protein, while smaller defects on flat membranes can inhibit the folding of the same protein domain^42^. This suggests that the size and nature of membrane defects play a critical role in modulating the structural behavior of IDPs upon binding. Furthermore, the depth distribution of membrane packing defects is crucial for understanding membrane-protein associations, as deeper defects can facilitate protein anchoring and subsequent binding^43^. The presence of defects in lipid membranes can act as attractive sites for proteins, with proteins being drawn to regions of lipid bilayers where hydrophobic defects are sensed^44^. These defects serve as key determinants for the specific membrane binding of peripheral proteins, highlighting their importance in cellular processes^45^. Moreover, the binding of IDPs to membranes is influenced by the membrane curvature. Proteins containing amphipathic helices, such as *α*S, sense membrane curvature through their interaction with strongly curved membranes, mediated by lipid packing defects^46^. The binding mechanism of these proteins to membranes is enhanced by positive feedback on lipid packing defects, leading to membrane deformation and potential fission^47^ what might underlie the cooperative type of interaction between *α*S and lipid membranes. Finally, the third parameter, depending on the model, is either interdigitation of the membrane or membrane thickness. In the case of interdigitation, the correlation is relatively small (from 0.003 to 0.06). With such a low amplitude of correlation, it is unlikely that this parameter has a significant contribution towards protein affinity to the membrane. Interestingly though, in the case of model #3 there is a strong negative correlation for the logarithmic value of membrane thickness. This could indicate that *α*S protein prefers membranes with greater membrane thickness. This would be in agreement with the fact that fluctuations in membrane thickness are often localized around defects, suggesting that the dynamics of these defects can lead to measurable changes in the overall membrane thickness.^48^ Furthermore, it has been postulated that the mechanism of defect formation and the structural properties of membranes are interlinked, with thicker membranes potentially being more prone to defects under certain conditions.^49^ Interestingly though, as we do not include curvature in our set of parameters, it is possible that presence of both membrane thickness and interdigitation is the effect of a lurking variable. After all, the presence of high membrane curvature can induce interdigitation of acyl chains, leading to a significantly broadened melting transition and increased disorder of the lipid fatty acyl chains, resulting in membrane thinning^50^. Finally, it should be noted that protein experiments are performed on highly curved LUVs while membrane characterization is performed on monolayers, GUVs and simulations of planar membranes. While determined properties of membranes are global properties, it should be noted that LUVs might have some of those properties locally altered (such as defects). This could contribute to a small mismatch between the behavior of the proteins due to slightly different membrane characteristics in LUVs vs GUVs and planar membranes.

## Material & Methods

Lipids POPC (1-palmitoyl-2-oleoyl-glycero-3-phosphocholine), C1P 18:1/d18:1 (N-oleoyl-ceramide-1-phosphate), C1P 16:0/d18:1 (N-palmitoyl-ceramide-1-phosphate), POPA (1-palmitoyl-2-oleoyl-sn-glycero-3-phosphate), DPPA (1,2-dipalmitoyl-sn-glycero-3-phosphate) and cholesterol were purchased from Avanti Polar Lipids (Alabaster, AL, USA). Fluorophore DilC was purchased from Thermo Fisher (International). The structures of lipids are presented in Figure S12 , the amino acid sequence of *α* -synuclein protein is presented in Figure S13 . HEPES was purchased from Carl Roth (International). The ultrapure water used in experiments was obtained from the Water Purification System (Milipore). Imidazole, TEMED (N,N,N’,N’-tetramethyl ethylenediamine), lysozyme from chicken egg white were purchased from Sigma-Aldrich (international). IPTG (isopropyl-*β* -D-thiogalactopyranosid), TRIS (tris-(hydroxymethyl)-amino methane), PMSF (phenylmethyl sulphonyl fluoride), Triton^®^ X-100, HEPES (N-2-hydroxyethylpiperazine-N’-2-ethane sulfonic acid) and Rotiphorese^®^ Gel 30 were purchased from Roth (international). Phusion^®^ High-Fidelity DNA Polymerase and DpnI enzyme were purchased from New England Biolabs (Ipswich, MS, USA). Ultrapure dNTP Mix, agarose electrophoresis grade, and OMNI nuclease were purchased from EURx (Poland). NucleoSpin^®^ plasmid isolation kit and the NucleoSpin^®^ gel and PCR cleaning kit were purchased from Macherey-Nagel (Germany). PierceTM protease inhibitor tablets without EDTA were purchased from Thermo Scientific (international). TALON^®^ Metal Affinity Resin was purchased from Takara Bio (Mountain View, CA, USA). Glycerol, sodium chloride and ammonium persulfate were purchased from POCH (Poland). LB Miller broth was purchased from IBI Scientific (international). Kanamycin sulfate was purchased from BioShop (Poland). Coomasie Brillat Blue R-250 was purchased from PanReac AppliChem (Germany). pET His6 GFP TEV LIC cloning vector (1GFP) was a gift from Scott Gradia (Addgene plasmid 29663 ; http://n2t.net/addgene:29663 ; RRID:Addgene_2_9663).

### Expression and purification of α-synuclein

Protein production and purification were carried out as described for HisTag-*α*S-mEGFP in Drabik et al.^51^. More details are provided in Supplementary Information Section 5.2 .

### Interaction analysis using Microscale Thermophoresis

MST experiments were conducted using the Monolith NT.115 Nano equipped with a blue excitation laser and green fluorescence filter, purchased from NanoTemper Technologies (GmbH, Munich, Germany). *α*S-SiriusGFP protein at a concentration of 30 nM was incubated with a series of dilutions of lipid vesicles with varying lipid compositions in HBS buffer (20 mM HEPES, 150 mM NaCl, pH 7.4). The samples were then transferred into Monolith Standard Capillaries from NanoTemper Technologies (GmbH, Munich, Germany) for measurement. MST measurements were performed using a blue excitation laser at 100% power and an infrared laser set to a high power level. Fluorescence changes upon activation of the infrared laser were observed for 5 seconds. To plot the binding curves, the fluorescence change in each capillary was analyzed 2.5 seconds after activation of the infrared laser. After plotting the relationship between the protein fluorescence change and the logarithm of lipid concentration, the data were fitted to the dose-response curve with variable Hill slope (equation 1) using OriginPro 2024b software, where A1 is the bottom asymptote, A2 is top asymptote, *LOG*_*x*0_ is center of the sigmoidal curve (*EC*_50_) and p is the Hill slope (Hill coefficient). This fitting yielded parameters characterizing protein binding to lipid vesicles (*EC*_10_/*EC*_20_/*EC*_50_/*EC*_80_/*EC*_90_ and the Hill coefficient).

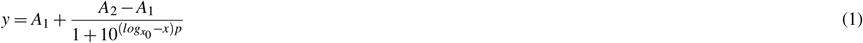

### Vesicle Preparation

Large unilamellar vesicles (LUVs) were prepared using a combination of freeze-thaw and extrusion methods. Briefly, the lipid mixtures in the solvent were dried under a nitrogen stream and placed in a vacuum desiccator overnight to eliminate any remaining organic solvents. The dried lipid films were rehydrated in HBS buffer (20 mM HEPES, 150 mM NaCl, pH 7.4) to reach a final lipid concentration of 5 mg/ml. The obtained multilamellar vesicles underwent 10 freeze-thaw cycles between liquid nitrogen and 45^*°*^C. This was followed by an extrusion procedure using Avanti mini-extruder. The size and zeta potential of the vesicles were evaluated using a ZetaSizer Nano ZS (Malvern Instruments, Malvern, UK) ( See Supplementary Information Section 5.3 for more information) .

Giant unilamellar vesicles (GUV) were obtained using a modified electroformation method^24^. Briefly, 10 µl of 1 mM POPC/C1P or POPC/Chol/C1P mixture and fluorescent probe mixture (0.5m%) in chloroform was distributed equally along the platinum electrodes and dried under vacuum overnight. The electrodes were then submerged in aqueous non-conductive solution and a sinusoidal 10 Hz AC electric field was applied for 4 h with 1 V increase for each hour.

### Molecular Dynamics Simulations

The full atomistic MD simulation was performed using NAMD 2.13^52^ software with CHARMM36 force fields^53^ under NPT conditions (constant: Number of particles, pressure, and temperature). Lipid membrane systems consisted of 648 lipid molecules (324 on each of the leaflets). The system was hydrated in such a way that 75 water molecules per lipid molecule were used. All systems were additionally neutralized with positive counter ions. All systems were equilibrated using the standard equilibration procedure. The total simulation time for all systems was at least 90 ns with the last 10 ns used for analysis. Simulations were carried out under 22^*°*^C (285.15K). C1P molecules force fields were taken from previous work.^24^ The systems are analyzed to obtain area per lipid, membrane thickness, bending rigidity, tilt rigidity, compressibility, lateral diffusion coefficient, interdigitation, order parameter, and acyl chain accessibility following the procedures established in our previous works^23^. A detailed description of the performed characterization is presented in Supporting Information section 5.4 .

### Surface PressureArea (π − A) Isotherm Measurement and Analysis

The *π* − *A* isotherms of C1P, POPC/C1P and POPC/Chol/C1P monolayers were measured using a computer-controlled Langmuir-type film system (Kibron). Before measurement, the trough (205 x 60 *mm*^2^) was carefully cleaned with water and ethanol. This was followed by filling the trough with 5 mM HEPES/KOH (7.4 pH) as the aqueous subphase. The lipid solution in hexane was spread over the subphase between the barriers. The concentration of lipids was determined using the total phosphorus assay method^54^. After the solvent was evaporated (about 10 min), the monolayer was compressed at a rate of 10 mm/min. The compression was carried out with two sliding barriers moving with the same speed from the edge of each compartment to its center, where the Wilhelmy plate, used as a surface pressure sensor, was placed. The subphase temperature was kept at 22.0±0.5^*°*^C. The measurements under the same conditions were repeated to obtain at least five *π* − *A* isotherm profiles. A monolayer was compressed until the surface pressure was 3 mN/m lower than the collapse pressure (which was determined during the first run). Both Area Per Lipid (APL) and excess free energy of mixing (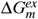) were analyzed at a pressure equal to 33 mN/m. The analysis was done as in our previous work.^23^ A more detailed description is provided in Supplementary Information Section 5.5 .

### Flicker Noise spectroscopy

The GUV series of images were recorded using Stellaris 8 confocal microscopy (Leica, Germany) equipped with an HC PL APO 86×/1.20 water immersion objective (Leica, Germany) with a pinhole size of 0.5 Airy units. 256x256 pixels images were recorded with a hybrid (HyD) detector with pixel size ranging from 0.07 µm to 0.12 µm with video integration time ranging from 148 ms up to 189 ms depending on the zoom magnitude. The samples were illuminated with a white laser set at 560 nm, emitted light was recorded from 570 to 630 nm. The series usually consisted of 1200 images. The images were analyzed to obtain the bending rigidity values as described in detail elsewhere^55^. The radii of the investigated vesicles ranged from 3.2 µm to 11.6 µm.

### Membrane generalized polarization measurement

To measure the generalized polarization of LUV vesicle membranes, vesicles were diluted with HBS buffer to a final total lipid concentration of 50 µM. Subsequently, C-Laurdan was added to the vesicle solutions to a final concentration of 100 nM, followed by a 30 minute incubation at room temperature. The emission spectra of the C-Laurdan probe embedded in the lipid vesicles, in the range of 300-540 nm, were recorded upon excitation at a wavelength of 340 nm. The measurements were performed using an FS5 spectrofluorometer (Edinburgh Instruments, UK) with excitation and emission slits set to 3 nm and an integration time of 0,1s / 1 nm at room temperature. Generalized polarization was calculated from the fluorescence spectra using Equation 2, where I is fluorescence intensity at a given wavelength.

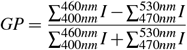

### Principle Component Analysis and Multiple Line Regression modeling

Principal component analysis was performed using OriginPro 2015 (OriginLabs) software. To determine models of cooperativity and effective concentrations on the physicochemical properties of membranes, multiple line regression (MLR) modeling was used. Specifically, each of the parameters determined using *in vitro* or *in vivo* studies was subjected to various mathematical operations (the list of the 81 parameters is presented in Supplementary Table S3 ). Either Hill Coefficient or *EC*_50_ values were modeled. The weights were calculated as 1/uncertainty of the modeled value. For membranes with two biphasic nature the lower value was taken. MLR model was trained on the results of all investigated membranes - C1P, POPC:C1P 8:2 and POPC:Chol:C1P 5:3:2. To increase the size of the training set, it was supplemented with results from PA membranes^23^. The threshold level of *R*^2^ of the model was set at 0.85. In case the model scored higher, it was used for validation. The validation consisted of two membranes - POPC:C1P (16:0) 9:1 and POPC:Chol:C1P (18:1) 6:3:1. To ensure the validity and reliability of the model, the sum of the residue of both training and validation (the difference between the measured and modeled values) was calculated. The models were sorted on the basis of descending order and three best were selected.

### Statistics

In order to test for the significant difference between the parameters describing protein - membrane interaction and membrane generalized polarization values, the Brown-Forsythe and Welch’s ANOVA tests were used with the significance level at 0.05 with the following Dunnett’s multiple comparisons test (*α* = 0.05) using the GraphPad Prism 10 software. For parameters describing the physicochemical characterization of C1P membrane systems, unless otherwise specified, the one-way ANOVA test was used with the significance level at 0.05. The Tukey test was used as a post hoc test. All statistical analyzes were performed with the OriginPro 2015 software (OriginLabs). Average values are presented with standard deviation.

## Conclusions

In this study, we presented an in-depth investigation of interactions between *α*-synuclein and lipid membranes enriched with ceramide-1-phosphate (C1P) and cholesterol. Our study addressed three novel issues. First, we postulated that C1P is a lipid molecule that interacts with *α*S. Indeed, our results confirm that interactions between those particles occur; however, they are subject to C1P concentration and acyl chain configuration in the investigated membrane. Membranes containing saturated C1P (C16:0) exhibited the highest binding affinity and cooperativity for *α*S, especially in the absence of cholesterol. In contrast, unsaturated C1P (C18:1) reduced binding affinity, with cholesterol modulating this interaction. The biphasic binding observed in the C1P (16:0) systems suggests multiple binding modes or sites, which may be attributed to the formation of domains in the membrane. The presence of cholesterol had different effects on the C1P sub-populations. Although cholesterol did not significantly affect the cooperativity of binding affinity in saturated C1P membranes, it enhanced the affinity of *α*S for unsaturated C1P membranes. This suggests that cholesterol may stabilize certain membrane properties that promote protein binding in specific lipid environments.

The second novel aspect is the biophysical characterization of C1P membranes. Numerous similarities between PA and C1P enriched membranes were found. One of such similarities was observed in a monolayer study in which a compression effect similar to the one in POPC:POPA systems was observed. The peculiar behavior of the lipid molecule C1P (16:0) was observed. When in homogeneous membranes, their effect on membrane properties was reversed when in mixtures with POPC or cholesterol. This effect was not observed in the case of C1P (18:1). C1P (16:0) was also a lipid that exhibited most difference in APL values between membrane and monolayer system, specifically (69.5 ± 2.4) Å^2^ and (61 ± 2) Å^2^, indicating that this lipid types dynamically adapt to given membrane physicochemical state.

The final novel issue is the approach to determine the relationship between the membrane’s physicochemical properties and protein-membrane binding cooperativity and affinity. In fact, structural parameters such as membrane thickness, compressibility, and lateral diffusion were found to correlate with *α*S binding. In the case of cooperativity, we observed strong dependency on the ordering of lipid molecules in the membrane. The negative dependency of bending rigidity implies that stabilization of the membrane curvature is a crucial factor. The higher mixing energy cost of the membrane system resulted in a reduced cooperativity of protein incorporation process. We postulate that this is related to the presence of heterogeneity, which contributes to the limited area of proteins to incorporate and the decrease of cooperativity due to surface limitation. In the case of affinity (for which *EC*_50_ is used as indicator) we observed strong correlation with global polarization, suggesting that domain formation does influence the incorporation of the investigated protein. Furthermore, the models obtained indicate a dependency on the presence of defects in the membrane, specifically the size of the defects. *α*S proteins tend to incorporate when the acyl chain region is sufficiently exposed (0.3 − 0.4*nm*), but not too exposed (0.5*nm*). Finally, a negative correlation with membrane thickness was observed, indicating that *α*S, as a small protein, prefers membranes with a smaller membrane thickness. These findings provide insights into the molecular mechanisms underlying protein-lipid interactions and could be relevant for understanding *α*S-related pathologies, such as Parkinson’s disease. Further studies could explore how these interactions affect *α*S aggregation and the formation of toxic oligomeric species in neurodegenerative diseases.

## Acknowledgements

This work was possible thanks to the financial support from the National Science Centre (Poland) grant no. 2018/30/E/NZ1/00099.

## Author contributions statement

Conceptualisation - AC DD PH, Resources - AC DD, Formal Analysis - DD PH, Simulations - DD, Protein expression - PH, Investigation - DD PH KC, Funding Acquisition - AC, Validation - DD PH, Methodology - DD PH, Visualization - DD PH, Project Administration - AC DD, Writing of original draft - DD PH, Review & Editing - all authors.

### Additional information

To include, in this order: **Accession codes** (where applicable); **Competing interests** (mandatory statement).

The corresponding author is responsible for submitting a competing interests statement on behalf of all authors of the paper. This statement must be included in the submitted article file.

