## Supporting Information for "Is efficiency of protein incorporations governed by membrane properties? An *α*-synuclein and ceramide-1-phosphate investigation"

### Supporting Information: Is efficiency of protein incorporations governed by membrane properties? An α-synucleine and ceramide-1-phosphate investigation.

Dominik Drabik[1,2,+], Piotr Hinc[1,+], Karolina Cierluk[1], and Aleksander Czogalla[1]

1 Department of Cytobiochemistry, Faculty of Biotechnology, University of Wroclaw, F. Joliot-Curie 14a, 50-383 Wrocław, Poland.

2 Department of Biomedical Engineering, Faculty of Fundamental Problems of Technology, Pl. Grunwaldzki 13, 50-377 Wrocław, Poland.

+ these authors contributed equally to this work

#### 1. Membrane characterization

##### 1.1. APL comparison between Monolayer and simulation study

In Figure S1 a comparison between obtained and theoretically calculated values from corresponding mono-lipid systems is presented. One can clearly see that there is slight discrepancy between the MD and monolayer obtained parameters – especially for systems with cholesterol. The most curious case can be observed in the case of POPC:Chol:C1P16:0, where theoretical values from both experiment and simulation are in agreement, but value from MD is lower and value from experiment is higher. We think that this is partially due to the compression effect that was observed, but also the effect of C1P on behaviour of POPC and Chol in the membrane. Similarly, like in the case of POPA [1], Results from experimental studies of membrane parameters are presented in Tab S1.

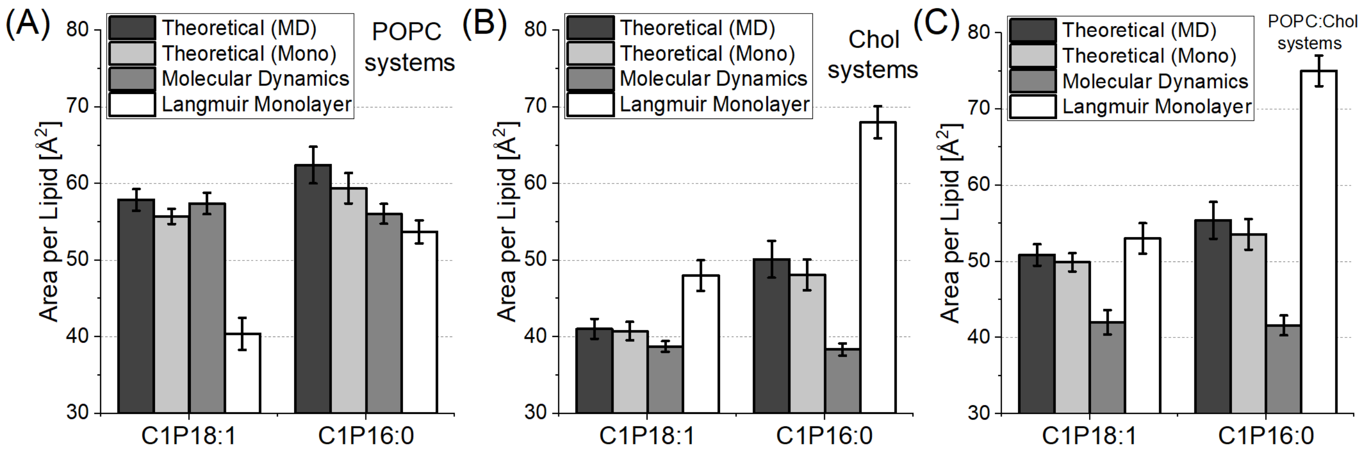


Figure S1. Comparison between values theoretically calculated from homogeneous systems and obtained from heterogeneous ones. Results are presented for both molecular dynamics and Langmuir monolayer studies. (A) Comparison between POPC:C1P systems; (B) comparison between Chol:C1P systems; (C) Comparison between POPC:Chol:C1P systems.

Tab S1. Membrane parameters obtained from experimental studies - Langmuir monolayers, Flicker Noise spectroscopy and GP studies.

| Membrane | Area per lipid [A^2] | Monolayer compressibility $C_s$ [mN/m] | Mixing parameter [$10^{-23}$ J] | Excess free energy of mixing $\Delta G_m^{ex}$ [J] | Bending rigidity κ [fold $K_BT$] | Bending rigidity κ [10^-19 J] | Number of vesicles for analysis | GeneralizedPolarisation GP |
|---|---|---|---|---|---|---|---|---|
| **C1P18** | 42.6 ± 1.0 | 67 ± 12 | - | - | - | - | - | - |
| **C1P16** | 61 ± 2 | 197 ± 44 | - | - | - | - | - | - |
| **POPC:C1P18** | 40.4 ± 2.1 | 60 ± 14 | -624.3 | -3760 | 150 ± 37 | 6,1 ± 1,5 | 14 | -0,099±0,002 |
| **POPC:C1P16** | 53.7 ± 1.5 | 55 ± 8 | 179.8 | 1080 | 39 ± 13 | 1,6 ± 0,5 | 10 | -0,038±0,009 |
| **Chol:C1P18** | 48 ± 2 | 122 ± 18 | 908.0 | 5470 | - | - | - | - |
| **Chol:C1P16** | 68.0 ± 2.1 | 192 ± 35 | 1047.4 | 6310 | - | - | - | - |
| **POPC:Chol:C1P18** | 53 ± 2 | 53 ± 6 | 742.1 | 4470 | 352 ± 98 | 14,4 ± 4,2 | 18 | 0,169±0,005 |
| **POPC:Chol:C1P16** | 75 ± 2 | 81 ± 18 | 1107.0 | 6670 | 313 ± 97 | 13 ± 4 | 12 | 0,272±0,004 |

##### 1.2. Compression effect observed in Monolayers

While the meaning of APL is straightforward, the free energy of mixing is a measure of energy change due to mixing – to consider a process spontaneous the energy of mixing should be negative, otherwise an external energy is required for the mixing of membranes. Obtained isotherms are presented in Figure S2.A. Similarly, like in the case of POPC:POPA [1], the compression effect of monolayers was observed as shown in the POPC:C1P18:1 system. The established energies of mixing are presented in Figure S2.B. The lipid ratio of investigated systems are C1P:PC 2:8, C1P:Chol:PC 2:3:5, Chol:C1P 4:6.

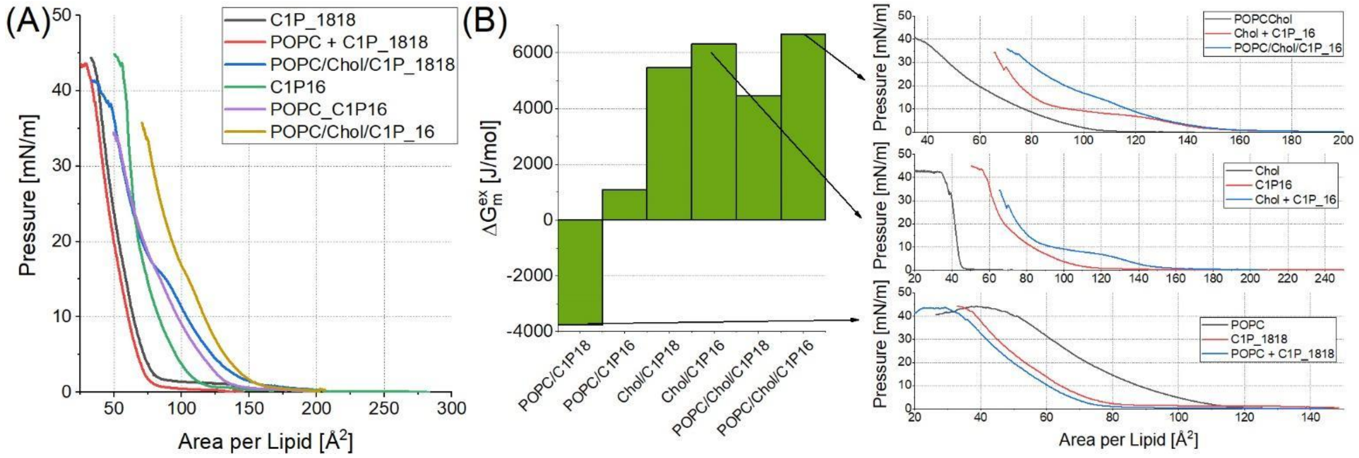


Figure S2. (A) Obtained isotherms of investigated monolayer systems. (B) Calculated energies of mixing with detailed visualisation of isotherms for three most extreme cases.

##### 1.3 Membrane Parameters obtained from MD

In this section parameters obtained from MD studies are presented in Tab S2. The detailed visualisation of APL Voronoi analysis is presented in Figure S3.

Tab S2. Membrane parameters obtained from simulation studies.

| Membrane | Membrane Thickness [nm] | Area per lipid [A^2] | Calculated bending rigidity κ [10^-19 J] | Calculated tilt rigidity $\kappa_{tilt}$ [10^-20 J] | Area compressibility $K_A$ [mN/m] | Lateral diffusion D [um2/s] | Interdigitation [A] | Scd sn1 | Scd sn2 |
|---|---|---|---|---|---|---|---|---|---|
| **C1P18** | 4.22 ± 0.03 | 46.8 ± 1.3 | 1.33 ± 0.03 | 4.03 ± 0.06 | 2754 ± 134 | 2.39 ± 0.04 | 5.05 ± 0.33 | 0.20 ± 0.09 | 0.28 ± 0.08 |
| **C1P16** | 2.92 ± 0.01 | 69.5 ± 2.4 | 3.92 ± 0.09 | 17.9 ± 0.3 | 7843 ± 173 | 0.55 ± 0.02 | 11.2 ± 0.2 | 0.35 ± 0.09 | 0.36 ± 0.10 |
| **POPC:C1P18** | 4.07 ± 0.02 | 57.4 ± 1.4 | 1.25 ± 0.02 | 3.82 ± 0.06 | 721 ± 49 | 3.65 ± 0.04 | 6.73 ± 0.42 | 0.13 ± 0.11 | 0.15 ± 0.12 |
| **POPC:C1P16** | 4.19 ± 0.03 | 56.1 ± 1.3 | 1.59 ± 0.03 | 4.53 ± 0.08 | 413 ± 61 | 2.33 ± 0.04 | 4.89 ± 0.25 | 0.24 ± 0.09 | 0.18 ± 0.11 |
| **Chol:C1P18** | 3.75 ± 0.01 | 38.8 ± 0.7 | 1.96 ± 0.05 | 6.3 ± 0.1 | 12532 ± 309 | 2.11 ± 0.09 | 3.40 ± 0.22 | 0.23 ± 0.07 | 0.31 ± 0.07 |
| **Chol:C1P16** | 3.86 ± 0.02 | 38.4 ± 0.8 | 1.87 ± 0.05 | 7.26 ± 0.07 | 16981 ± 359 | 0.43 ± 0.02 | 2.9 ± 0.3 | 0.32 ± 0.07 | 0.33 ± 0.06 |
| **POPC:Chol:C1P18** | 4.12 ± 0.02 | 42.0 ± 1.6 | 2.23 ± 0.05 | 10.2 ± 0.02 | 6617 ± 175 | 2.48 ± 0.08 | 3.55 ± 0.24 | 0.23 ± 0.07 | 0.28 ± 0.08 |
| **POPC:Chol:C1P16** | 4.18 ± 0.02 | 41.6 ± 1.3 | 2.61 ± 0.06 | 12.5 ± 0.2 | 7031 ± 173 | 1.09 ± 0.03 | 3.72 ± 0.32 | 0.33 ± 0.07 | 0.31 ± 0.07 |

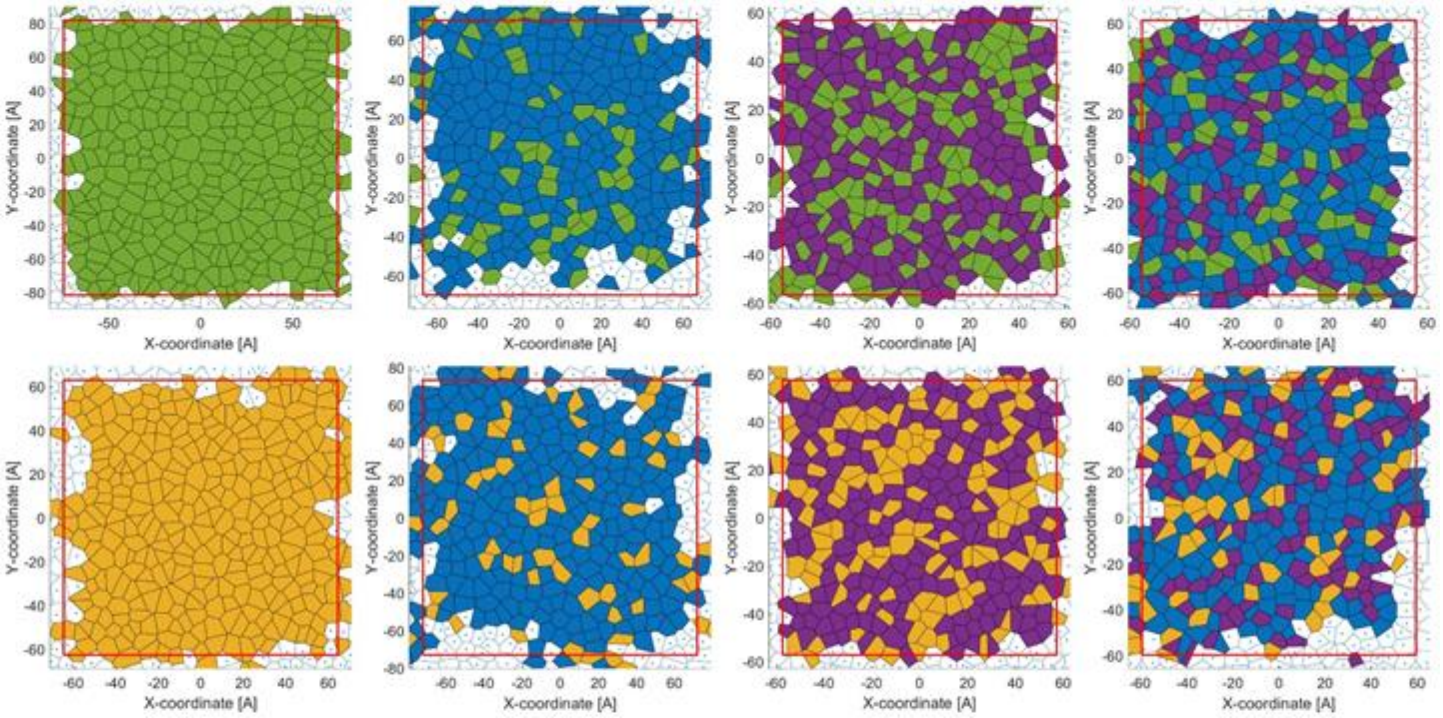


Figure S3. Visualisation of Voronoi tessellation of lipids for each of the investigated systems. The upper row presents systems with C1P16:0, the lower row presents systems with C1P18:1. From left to right: C1P, POPC:C1P, Chol:C1P and POPC:Chol:C1P systems. The color legend is as follows: Green-C1P16:0, Blue-POPC, Violet-Chol, Orange-C1P18:1.

##### 1.4 Flicker noise and GP additional information

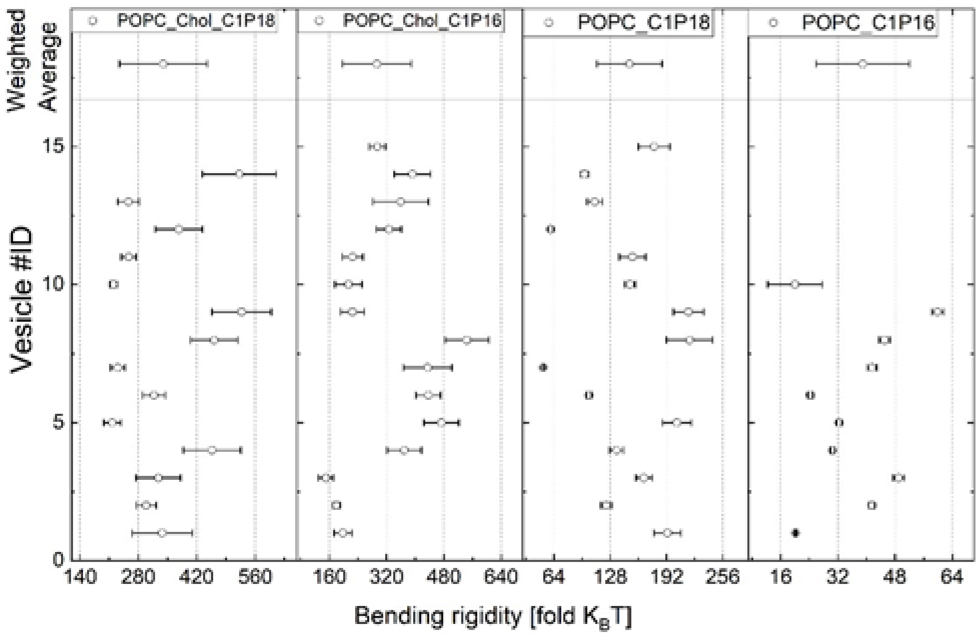


Fig S4. Individual bending rigidity determination for each of the investigated vesicles. The weighted average value is presented in top part of the plot.

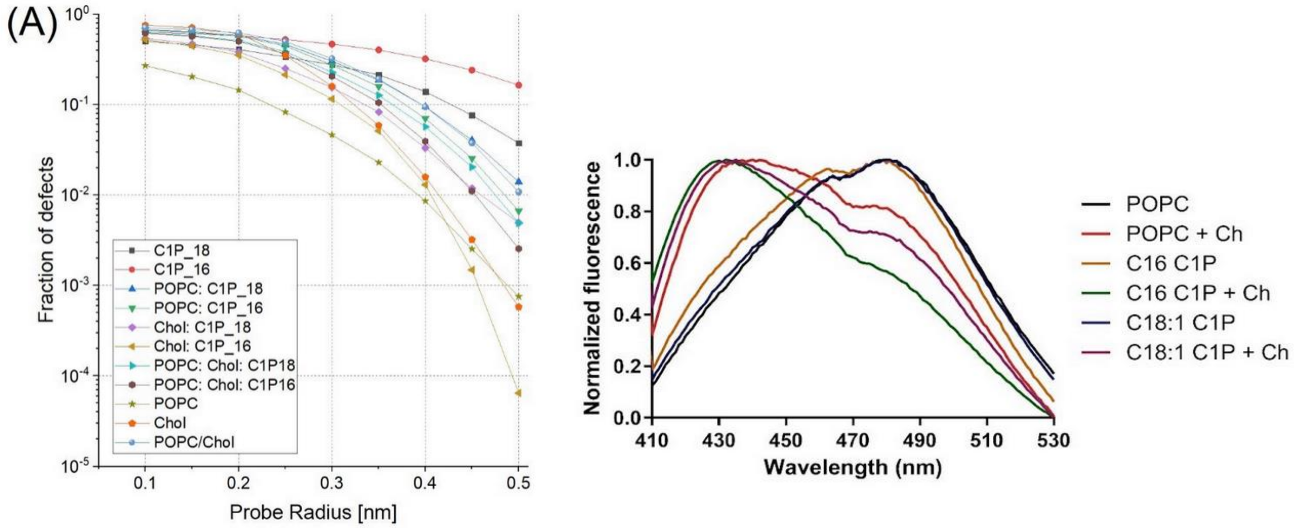


Fig, S5. (A) Calculated fraction of defects for investigated systems using molecular dynamics study. (B) Averaged fluorescence emission spectra of C-Laurdan probe embedded into studied membrane systems.

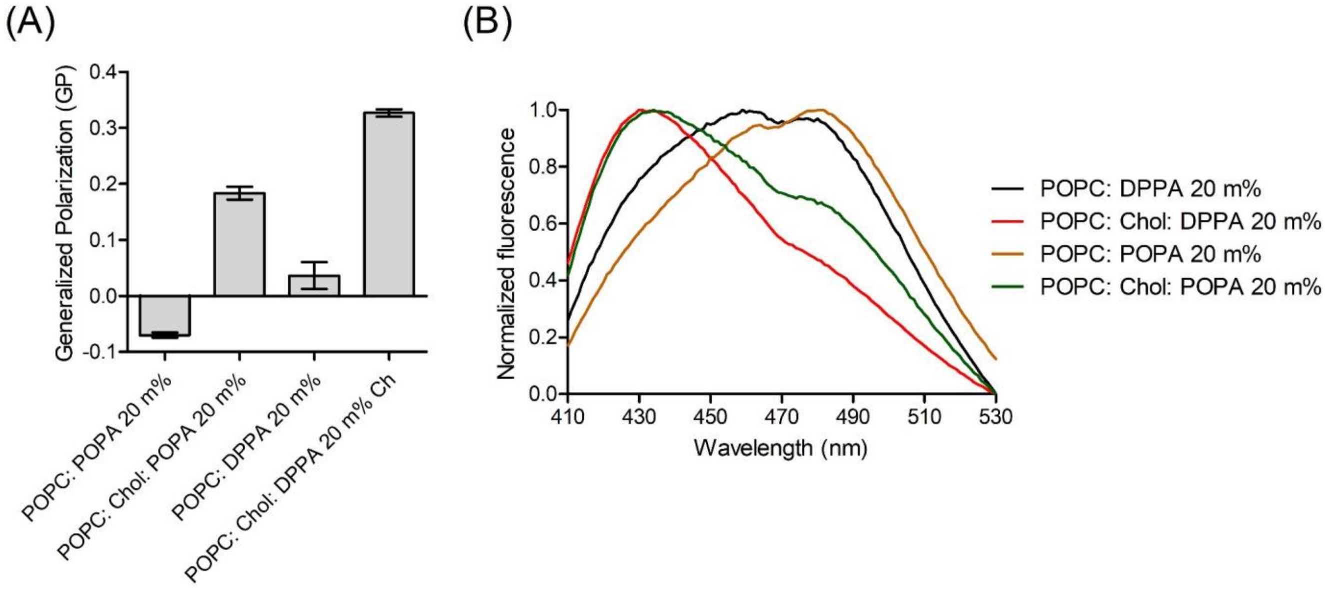


Fig S6. Calculated GP values for systems containing phosphatidic acid used as a part training data set (A) and corresponding averaged fluorescence emission spectra of C-Laurdan probe embedded into those systems (B).

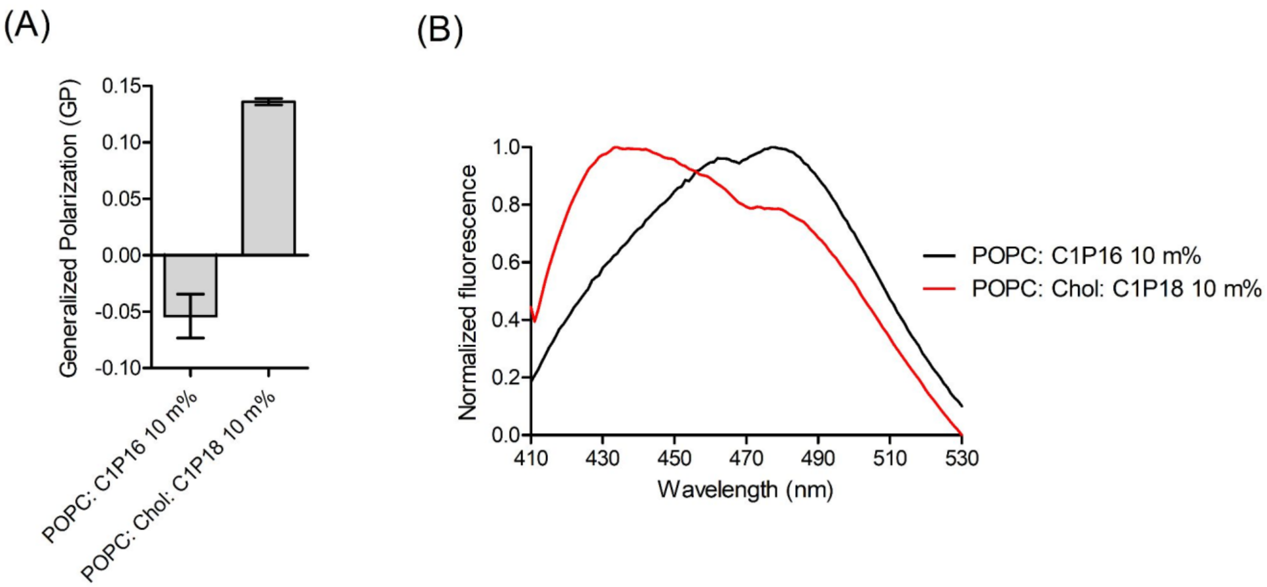


Fig S7. Calculated GP values for systems used as validation data set (A) and corresponding averaged fluorescence emission spectra of C-Laurdan probe embedded into those systems (B).

##### 3 Interaction of α-synuclein with membranes containing both C1P and PA

The results of $EC_{50}$ and Hill coefficients for all membrane systems used in MLR modelling are presented in Table S3. Additionally binding curves of α-synuclein - lipid vesicles interactions for systems containing phosphatidic acid and systems with C1P used as validation data set are presented in Figure S8 and Figure S9 respectively.

Tab S3. Binding parameter ($EC_{50}$ and Hill coefficient), obtained from fitting of binding curves to dose-response model, for all systems used for analysis of protein - membrane interaction. Data are shown as parameter value ± SE.

| Vesicles lipid composition | $EC_{50}$ [μM] | | Hill coefficient | |
|---|---|---|---|---|
| POPC | 348.91 ± 19.51 | | 2.79 ± 0.44 | |
| POPC: Chol | 392.82 ± 43.27 | | 2.80 ± 0.79 | |
| POPC: 20 m% POPA | 28.61 ± 4.41 | | 0.79 ± 0.08 | |
| POPC: Chol: 20 m% POPA | 23.73 ± 1.99 | | 1.13 ± 0.10 | |
| POPC: 20 m% C1P18:1 | 1130.00 ± 583.09 | | 0.48 ± 0.05 | |
| POPC: Chol: 20 m% C1P18:1 | 49.37 ± 9.57 | | 0.80 ± 0.14 | |
| POPC: 20 m% DPPA | 816.03 ± 195.56 | 0.66 ± 0.16 | 1.56 ± 0.38 | 2.61 ± 1.44 |
| POPC: Chol: 20 m% DPPA | 1350.00 ± 108.26 | 11.41 ± 8.86 | 2.01 ± 0.30 | 0.79 ± 0.26 |
| POPC: 20 m% C1P16:0 | 241.18 ± 32.56 | 1.47 ± 0.06 | 2.24 ± 0.54 | 2.20 ± 0.26 |
| POPC: Chol: 20 m% C1P16:0 | 350.73 ± 72.39 | 1.25 ± 0.25 | 1.70 ± 0.71 | 2.61 ± 0.85 |
| POPC: 10 m% C1P16:0 | 2140.00 ± 344.83 | | 2.60 ± 1.17 | |
| POPC: Chol: 10 m% C1PC18:1 | 135.86 ± 62.01 | | 0.66 ± 0.15 | |

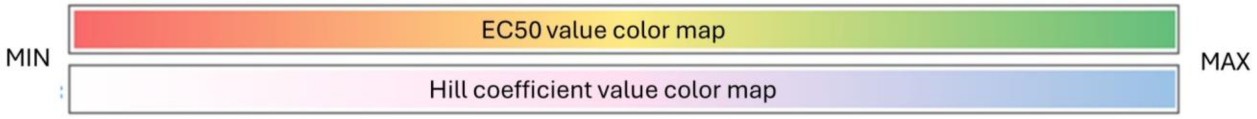

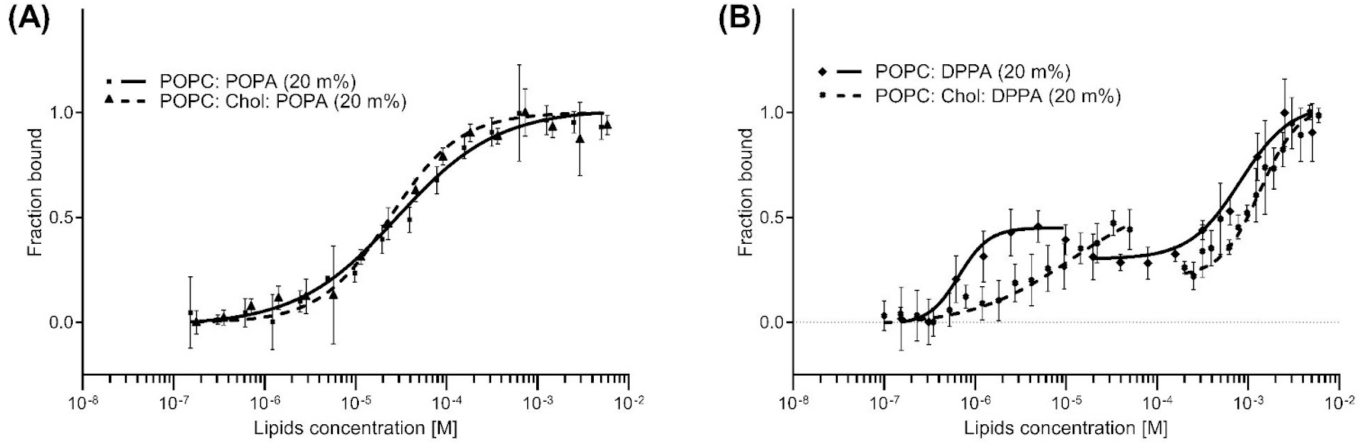


Fig S8. Binding curves of α-synuclein - lipid vesicles interactions for systems containing phosphatidic acid used as a part training data set.

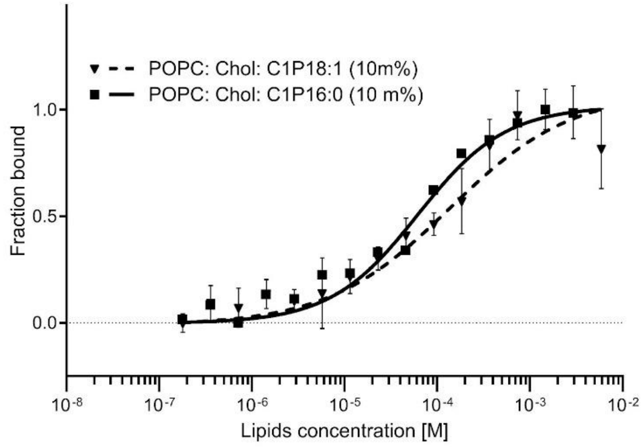


Fig S9. Binding curves of α-synuclein - lipid vesicles interactions for systems used as a validation data set.

In the case of membranes containing 20 m% saturated phosphatidic acid (16:0-16:0), a biphasic nature of binding was observed (Fig. S8B). The first phase, with an $EC_{50}$ of 1.56 ± 0.38 µM and a Hill coefficient of 2.61 ± 1.44, demonstrated a high-affinity interaction with strong positive cooperativity. The second binding phase showed an $EC_{50}$ of 816.03 ± 195.56 µM and a Hill coefficient of 0.66 ± 0.16, indicating a low-affinity binding site with negative cooperativity (Tab. S3).

For membranes containing 20 m% saturated PA and 30 mol% cholesterol, the binding was also biphasic (Fig. S8B). The first phase exhibited an $EC_{50}$ of 11.41 ± 8.86 µM and a Hill coefficient of 0.79 ± 0.26, suggesting a high affinity and slightly negative cooperativity of binding. The second binding phase had an $EC_{50}$ of 1350.00 ± 108.26 µM with a Hill coefficient of 2.01 ± 0.30, indicating low affinity with positive cooperativity (Tab. S3).

Membranes containing 20 m% unsaturated phosphatidic acid (16:0-18:1) displayed monophasic binding (Fig. S8A), characterised by an $EC_{50}$ of 28.61 ± 4.41 µM and a Hill coefficient of 0.79 ± 0.08. This low Hill coefficient suggests a lack of cooperativity or negative cooperativity. When 30 m% cholesterol was added to these membranes, the binding affinity has not changed significantly, as reflected in a $EC_{50}$ value of 23.73 ± 1.99 µM. The Hill coefficient increased to 1.13 ± 0.1, indicating a modest increase in cooperativity, however it is not statistically significant change (Tab. S3).

In membranes containing 10 m% saturated C1P (16:0), the binding was monophasic (Fig. S9) with an $EC_{50}$ of (2140.00 ± 344.83) µM and a Hill coefficient of (2.60 ± 1.17) (Tab. S3). Interestingly, there is much lower affinity compared to both modes/sites of binding compared to the system with twice the concentration of C1P ($p<0.0001$), the Hill coefficient indicated positive cooperativity with no significant difference compared to the corresponding system with higher concentration of C1P.

Similarly, in membranes containing 10 mol% unsaturated C1P (18:1) and 30 mol% cholesterol, the binding was monophasic (Fig. S9) with an $EC_{50}$ of 135.86 ± 62.01 µM and a Hill coefficient of 0.66 ± 0.15 (Tab. S3), indicating moderate affinity and non-cooperative or slightly negative cooperative binding showing no statistically significant difference in both values compared to corresponding system with higher concentration of C1P.

**4. Details of SAR, along with $EC_{10}$ $EC_{20}$ $EC_{80}$ and $EC_{90}$**

Tab S4. List of descriptors used in MLR study. The most left column is the determined parameter, the rest in the row are mathematically recalculated parameters. The first letter e indicates experimentally obtained parameter, while d indicates parameter from simulation.

| | | | | |
|---|---|---|---|---|
| e.kappa | sqrt(e.kappa) | (e.kappa)^2 | log(e.kappa) | - |
| e.APL | (e.APL)^2 | log(e.APL) | sqrt(e.APL) | - |
| e.Cs | (e.Cs)^2 | log(e.Cs) | sqrt(e.Cs) | - |
| e.dG | sign(e.dG) | sign(e.dG) * sqrt(\|e.dG\|) | log(\|e.dG\|) | - |
| e.GP | (e.GP)^2 | sign(e.GP)*sqrt(\|(e.GP)\|) | sign(e.GP)*log\|e.GP\| | exp(e.GP) |
| d.FP | - | d.Defects(0.1) | exp(d.Defects(0.1)) | - |
| d.Defects(0.2) | d.Defects(0.2)^2 | - | d.Defects(0.3) | sqrt(d.Defects(0.3)) |
| d.Defects(0.4) | sqrt3(d.Defects(0.1)) | - | d.Defects(0.5) | log(d.Defects(0.5)) |
| d,MT | (d,MT)^2 | log(d,MT) | sqrt(d,MT) | - |
| d.APL | log(d.APL) | sqrt(d.APL) | - | - |
| d.kappa | log(d.kappa) | sqrt(d.kappa) | - | - |
| d.tilt | log(d.tilt) | sqrt(d.tilt) | sqrt(d.tilt) | - |
| d.KA | log(d.KA) | sqrt(d.KA) | sqrt3(d.KA) | - |
| d.Intrdgt | exp(d.Intrdgt) | sqrt(d.Intrdgt) | (d.Intrdgt)^2 | sqrt3(d.Intrdgt) |
| d.Scd_sn1 | (d.Scd_sn1)^2 | sqrt(d.Scd_sn1) | - | - |
| d.Scd_sn2 | (d.Scd_sn2)^2 | sqrt(d.Scd_sn2) | - | - |
| d.max(H_up) | (d.max(H_up))^2 | (d.max(H_up))^3 | sqrt(d.max(H_up)) | exp(d.max(H_up)) |
| d.K_up | (d.K_up)^2 | (d.K_up)^3 | sqrt(d.K_up) | exp(d.K_up) |
| d.max(H_down) | (d.max(H_down))^2 | (d.max(H_down))^3 | sqrt(d.max(H_down)) | exp(d.max(H_down)) |
| d.K_down | (d.K_down)^2 | (d.K_down)^3 | sqrt(d.K_down) | exp(d.K_down) |

Abbevirations: kappa - bending rigidity coefficient; APL - area per lipid; Cs - monolayer compressibility; dG - mixing constant; GP - global polarization; FP - polar fraction in defect analysis; MT - membrane thickness; KA - compressibility modulus; Intrdgt - interdigitation; Scd - order parameter; H - mean curvature; K - gaussian curvature.

Two latter models of Hill Coefficient modelling are presented in Figure S.C. The resulting equations of those models are 1.1 and 1.2, respectively.

$$Hill = -1.00 * \log(e.kappa) - 0.561 * \log(|e.dG|) + 9.974 * d.Scd\_sn1 + 2.713 \quad (1.1)$$

$$Hill = -0.092 * \sqrt{e.kappa} - 0.543 * \log(|e.dG|) + 9.581 * \sqrt{d.Scd\_sn1} \quad (1.2)$$

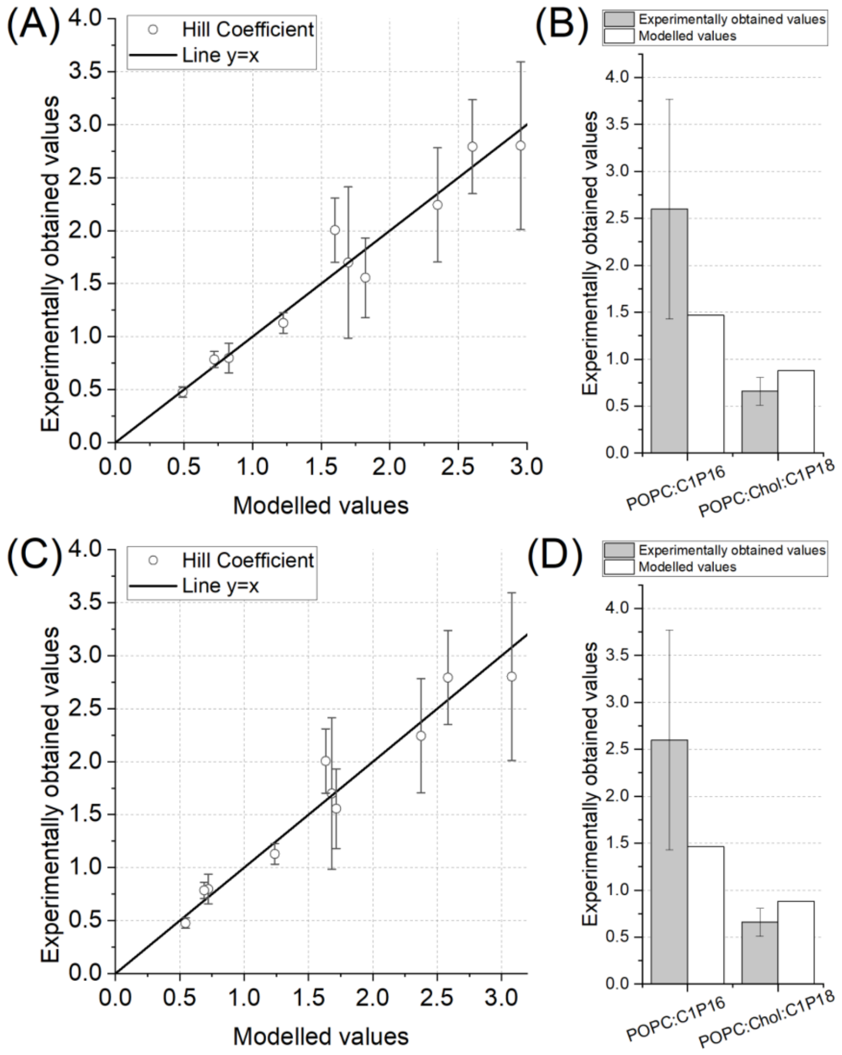


Fig S10. (A) The juxtaposition of experimentally determined values of Hill Coefficient (OY axis) and modelled values (OX axis) for #2 model. The black line represents perfect agreement between two datasets. (B) Comparison between experimentally obtained values of Hill coefficient (grey) and modelled values (white) in the training set for #2 model. (C) The juxtaposition of experimentally determined values of Hill Coefficient (OY axis) and modelled values (OX axis) for #3 model. The black line represents perfect agreement between two datasets. (D) Comparison between experimentally obtained values of Hill coefficient (grey) and modelled values (white) in the training set for #3 model.

Two latter models of $EC_{50}$ modelling are presented in Figure S.D. The resulting equations of those models are 1.3 and 1.4, respectively.

$$EC_{50} = -22.269 * d.Defects\_0.3 - 4.910 * \log(d.Defects\_0.5) + 0.174 * (d.Interdigit)^3 - 22.269 \quad (1.3)$$

$$EC_{50} = 41.148 * sqrt3(d.Defects\_0.4) - 7.579 * \log(d.Defects\_0.5) - 41.923 * sqrt(d.MT) - 15.434 \quad (1.4)$$

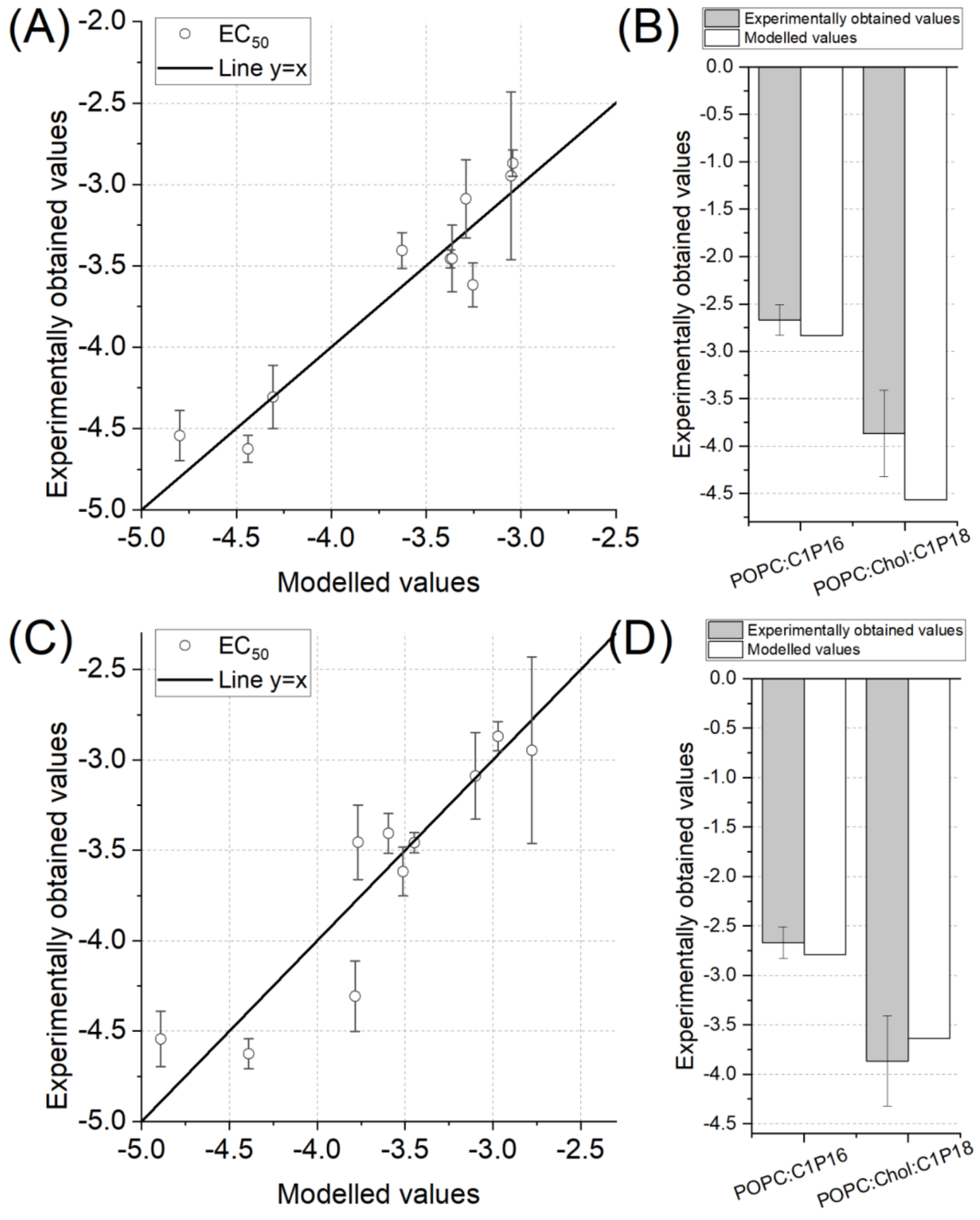


Fig S11. (A) The juxtaposition of experimentally determined values of $EC_{50}$ (OY axis) and modelled values (OX axis) for #2 model. The black line represents perfect agreement between two datasets. (B) Comparison between experimentally obtained values of $EC_{50}$ (grey) and modelled values (white) in the training set for #2 model. (C) The juxtaposition of experimentally determined values of $EC_{50}$ (OY axis) and modelled values (OX axis) for #3 model. The black line represents perfect agreement between two datasets. (D) Comparison between experimentally obtained values of $EC_{50}$ (grey) and modelled values (white) in the training set for #3 model.

#### 5 Materials and methodology section expansion

##### 5.1. Lipid structures

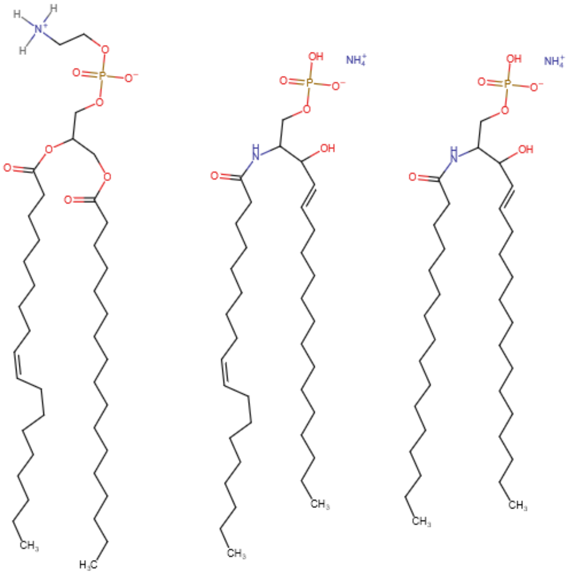

Figure S12. Chemical structure of investigated lipids in this study.

```
        10         20         30         40         50         60
MGHHHHHHHH GSGENLYFQG SMDVFMKGLS KAKEGVVAAA EKTKQGVAEA AGKTKEGVLY

        70         80         90        100        110        120
VGSKTKEGVV HGVTTVAEKT KEQVTNVGGA VVTGVTAVAQ KTVEGAGNIA AATGFVKKDQ

       130        140        150        160        170        180
MGKGEEGYPQ EGILEDMPVD PGSEAYEMPS EEGYQDYEPE AILEVLFQGP GVSKGEELFT

       190        200        210        220        230        240
GVVPILVELD GDVNGHKFSV RGEGEGDATN GKLTLKFICT TGKLPVPWPT LVTTLTYGVQ

       250        260        270        280        290        300
CFSRYPDHMK QHDFFKSAMP EGYVQERTIS FKDDGTYKTR AEVKFEGDTL VNRIELKGID

       310        320        330        340        350        360
FKEDGNILGH KLEYNFNSHN VYITADKQKN GIKANFKIRH NVEDGSVQLA DHYQQNTPIG

       370        380        390        400
DGPVLLPDNH YLSTQSKLSK DPNEKRDHMV LLEFVTAAGI TLGMDELYK
```

Figure S13. Amino acid sequence of produced 8×His - tagged murine α-synuclein fused with SiriusGFP. Yellow - HisTag; Blue - TEV protease cleavage site; Red - murine α-synuclein; Purple - HRV 3C protease cleavage site; Green - SiriusGFP

##### 5.2. More detailed Expression and purification of α-synuclein

To obtain plasmid construct for overexpression (in bacterial system) of 8×His - tagged murine α-synuclein fused with SiriusGFP (monomeric green fluorescent protein with greatly enhanced photostability [2] to avoid artefacts in MST experiments due to photobleaching) the plasmid encoding HisTag-aSyn-mEGFP, prepared for previously published studies [3], was subjected to a restriction-free cloning procedure to replace the mEGFP coding sequence with SiriusGFP. The Restriction Free Cloning procedure was performed according to the instruction described by van den Ent & Löwe [4] using the primers designed in https://www.rf-cloning.org/tool

- forward primer: AGTGCTGTTTCAGGGCCCGGGCGTGAGCAAGGGCGAGGAG
- reverse primer: GGTGGTGCTCGAGTCACATATGTCACTTGTACAGCTCGTCCATGC

To amplify SiriusGFP coding sequence, the pET His6 GFP TEV LIC cloning vector (1GFP) (addgene plasmid #29663) was used as a template. To confirm and validate performed cloning, the obtained DNA constructs were subjected to Sanger sequencing (Microsynth Seqlab GmbH, Germany).

Production of recombinant HisTag-α-synuclein-SiriusGFP protein was performed in *Escherichia coli* NiCo21 (DE3) strain. 200 milliliters of 50 µg/mL kanamycin-supplemented LB Miller broth was inoculated with overnight preculture and incubated at 37°C at 200 rpm agitation to reach culture optical density (λ = 600 nm) of approximately 0.7. Then the protein overexpression was induced by adding IPTG at a final concentration of 100 µM. Overexpression was carried out at 18°C at 180 rpm agitation for 18 h. Next, cells were harvested by centrifugation (10,000 × g, 20 min, 4°C) and lysed by re-suspending the pellet in 10 mL of ice-cold lysis buffer (10 mM HEPES, 500 mM NaCl, 1 mg/mL lysozyme, 0.5% Triton X-100, 25 U/mL OMNI Nuclease, 10 mM imidazole, 1 mM PMSF, 1× Price Protease Inhibitor Tablets EDTA free [pH 8.0]) and subsequent incubation for 1 h at 4°C under gentle mixing. Afterward, the suspensions were sonicated on ice for 15 min at 80% amplitude and 0.5 cycle (Hielscher UP100H Ultrasonic Processor with MS3 sonotrode). Lysates were then clarified by centrifugation (30,000 × g, 30 min, 4°C) and the supernatants were incubated with 1 mL of previously equilibrated TALON Metal Affinity Resin for 2 h at 4°C under gentle mixing. Then resin was packed into a chromatography column and washed with 100 mL of Wash Buffer 1 (10 mM HEPES, 500 mM NaCl, 10 mM imidazole, 20% glycerol [pH 8.0]), 100 mL of Wash Buffer 2 (10 mM HEPES, 300 mM NaCl, 10 mM imidazole, 20% glycerol [pH 8.0]), and 100 mL of Wash Buffer 3 (10 mM HEPES, 300 mM NaCl, 10 mM imidazole [pH 8.0]) until the absorbance (at λ = 280 nm) of flow-through buffer, measured in a 1-cm optical path quartz cuvette, decreased below 0.01. Then protein was eluted from resin with an elution buffer (10 mM HEPES, 300 mM NaCl, 200 mM imidazole [pH 8.0]). Eluted fractions were then dialyzed against 20 mM HEPES + 150 mM NaCl buffer (pH 7.4) and the protein concentration was determined spectrophotometrically (at λ = 280 nm) (Cary 1E UV-Visible Spectrophotometer) employing excitation coefficient calculated using the ProtParam tool (web.expasy.org/protparam). The purity and molecular weight of the produced protein was determined by SDS-PAGE in a Laemmli system [5] using Any kD™ Mini-PROTEAN® TGX™ Precast Protein Gels (BioRad Laboratories, USA) with the following Coomassie Brilliant Blue R-250 staining. The SDS-PAGE analysis is shown in Figure S14. The purified protein was then aliquoted, flash frozen in liquid nitrogen, and stored at −80°C.

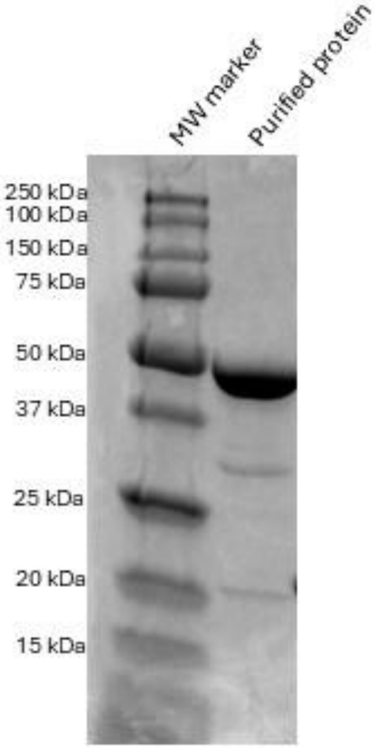


Figure S14. SDS-PAGE analysis of purified HisTag-α-synuclein-Sirius GFP protein.

**5.3. Vesicle Preparation**

For large unilamellar vesicles (LUVs) preparation lipids concentrations dissolved in organic solvents were measured through total phosphorus assay, while cholesterol levels were determined using a BioSystems Cholesterol Kit. Lipid mixtures were prepared in molar ratios of either 90 mol% POPC and 10 mol% C1P; 60 mol% POPC, 30 mol% cholesterol and 10 mol% C1P; POPC and 20 mol% PA/C1P; 50 mol% POPC, 30 mol% cholesterol and 20 mol% PA/C1P. For LUV preparation, the lipid mixtures (containing 5 mg of total lipids) in chloroform (or chloroform/methanol//water/ammonium hydroxide for DPPA and C16:0 C1P) were dried in a round-bottom flask under a nitrogen stream, then placed in a vacuum desiccator overnight to eliminate any remaining organic solvents. The dried lipid films were rehydrated in HBS buffer (20 mM HEPES, 150 mM NaCl, pH 7.4) to reach a final lipid concentration of 5 mg/ml. Obtained multilamellar vesicles underwent 10 freeze-thaw cycles between liquid nitrogen and 45°C. The vesicle suspension was then extruded sequentially using Avanti mini-extruder through polycarbonate membrane filters with pore sizes of 0.2 µm for 25 times, 0.1 µm for 25 times and through 0,05 µm for 15 times. (Whatman, United Kingdom). Extrusion was performed at a temperature of 35°C (for liposomes containing POPA and C18:1 C1P) or 70°C (for liposomes containing DPPA and C16:0 C1P). The size and zeta potential of the vesicles were evaluated using a ZetaSizer Nano ZS (Malvern Instruments, Malvern, UK) ( as shown in Figure S15). Measurements were performed at 25°C in 5 mM NaCl solution. For zeta potential measurement Zetasizer Nano Series Dip Cell Kit was used (Malvern Instruments, Malvern, UK)

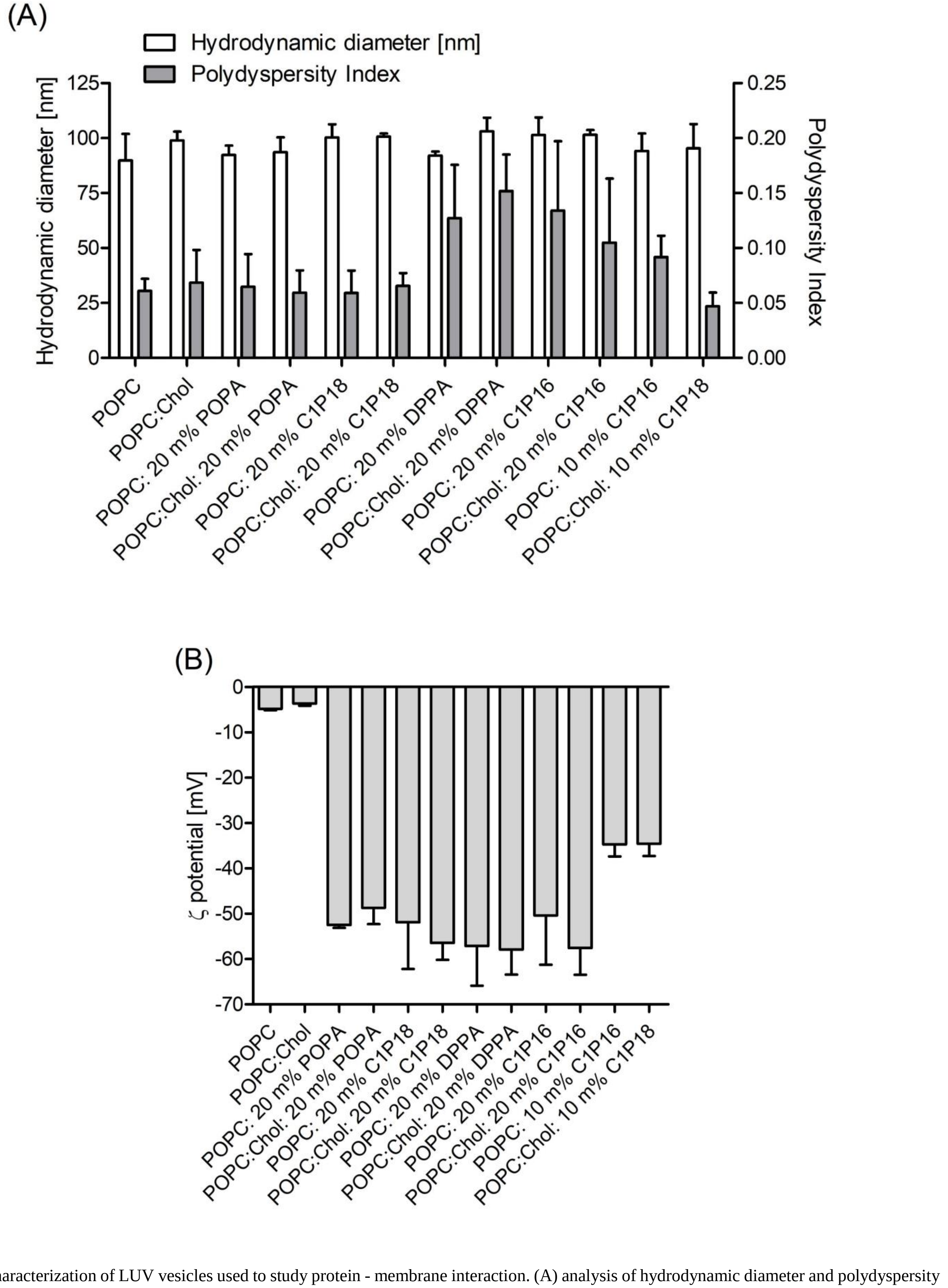


Figure S15**.** Characterization of LUV vesicles used to study protein - membrane interaction. (A) analysis of hydrodynamic diameter and polydyspersity index via dynamic light scattering; (B) analysis of zeta potential via electrophoretic light scattering.

Giant unilamellar vesicles (GUVs) were obtained using a modified method of model membrane formation for giant unilamellar vesicles (GUV) was used. Briefly, 10 µl of 1 mM POPC/C1P or POPC/Chol/C1P mixture and fluorescent probe mixture (0.5m%) in chloroform was distributed equally along the platinum electrodes and dried under the vacuum overnight. The electrodes were then submerged in aqueous non-conductive solution and a sinusoidal 10 Hz AC electric field was applied for 2 h with 1V voltage in a custom PTFE (polytetrafluoroethylene) electro-formation chamber [1]. However, for systems with cholesterol and C1P usage of such protocol resulted in formation of lipid droplet instead of lipid vesicles. Hence, prolonged 4 hour protocol [6] was used for those systems. Specifically, a square 1 Hz AC electric field was applied for 4 h with 1V increase for each hour.

##### 5.4. Force field modification and *in silico* membrane system characteristics

**Area per Lipid and Membrane Thickness.** Both membrane Area per Lipid (APL) and membrane thickness (MT) were determined using custom MATLAB script. Midpoint position between P and C2 atoms was used as a point position for each lipid. This was followed by Voronoi tessellation to obtain individual APL for each lipid molecule in every simulation time step. The APL value for whole membrane is determined by Gaussian fitting to histogrammed APL values of all molecules. The APL value for given type of molecule is determined by averaging. MT was calculated as an difference between mean height values of atoms in opposite leaflets. Parameter was determined for P, C1 and C2 atoms. For cholesterol corresponding C3, C2 and C1 atoms were selected.

**Bending rigidity and Tilt rigidity.** To determine mechanical parameters (focusing on bending rigidity coefficient, but also tilt modulus) a real-space fluctuation (RSF) method was used [7]. Specifically, a probability distribution for both tilt and splay is determined for all lipids over all analyzed time steps. Tilt θ is defined as an angle between the lipid director (vector between lipid head – midpoint between C2 and P atoms – and lipid tail – midpoint between last carbon atoms) and bilayer normal. Lipid splay $S_r$ is defined as divergence of an angle formed by the directors of neighboring lipids providing that they are weakly correlated.

**Compressibility.** Methods described by Doktorova *et al* [1, 8] was used to determine compressibility modulus of lipid membranes. Briefly, fluctuations of membrane thickness are used as a mean to determine leaflet compressibility modulus $K_A$.

**Lateral Diffusion Coefficient.** The Diffusion Coefficient Tool plugin was used for lipid molecules lateral (2D) diffusion determination [9]. It is calculated using Einstein's relation with mean square displacement (MSD) of the chosen molecular species.

**Interdigitation and Order Parameter.** Interdigitation and order parameters were determined using MEMBPLUGIN [10]. The obtained interdigitation parameter is defined as a width of the region of mass overlap. The obtained order parameters are used for analysis as average over one tail (sn1 or sn2).

**Acyl chain accessibility.** A protocol implemented by Boyd *et al* [11] was used for acyl chain accessibility determination. Briefly, the occupancy of each bilayer atom (from lowest to highest) is calculated based on Van der Waals radius. If it is an headgroup atom the sphere is marked as polar, otherwise as tail. Headgroup is defined as all atoms down to and including C2 atom. This is followed by dividing the simulation box into square grids with a grid spacing of 0.5 A. For each grid position a value of occupancy graph is checked. As a results a 2D map of either head or tail region is obtained, that is used to determine fraction of the bilayer surface allowing to access acyl chains region. This is followed by sphere probe analysis in order to ignore small gaps in headgroup coverage. For each point a sphere probe of radius ranging from 1nm up to 5nm is set. If the all grid points in the range of this probe are classified as tails, those grid points are assigned as defects. This way only hydrophobic patches with given sized and shapes are classified as true defects.

**Mean and Gaussian Curvature.** Membrane curvature was obtained using the MDAnalysis package [12], a Python library utilised in molecular dynamics simulations. MembraneCurvature module within MDAnalysis was employed, which allowed to calculate the mean curvature and Gaussian curvature based on a selection of reference atoms (in this case the z position of phosphorus atoms within each leaflet of the membrane).

##### 5.5 Details of Langmuir Monolayer Analysis

Area Per Lipid (APL) of investigated monolayers was collected at pressure equal to 33 mN/m. The area compressibility was calculated according to equation 1, where A is the area per molecule at the indicated surface pressure and π is the corresponding surface pressure [13].

$$C_s = \left(-\frac{1}{A}\right)\frac{dA}{d\pi} \qquad (1)$$

The excess free energy of mixing ($\Delta G_m^{ex}$) was calculated for π-A isotherms for pure and mixed monolayers following established protocol [1, 14]. The values were calculated at 33 mN/m. Briefly, the theoretical mean APL for non-interacting molecules was calculated as in equation 2, where $A_i$ - the mean molecular area, $X_1$, $X_2$ - mole fraction of the component 1 or 2, $A_1$ and $A_2$ - the mean molecular areas of pure components 1 or 2. If three component monolayers were analyzed an additional compartment was added.

$$A_i = X_1A_1 + X_2A_2 \ (2)$$

The mixing parameter was calculated according to equation 3 (in the case of two-compartment monolayers) and equation 4 (in the case of three-component monolayers). Calculations were carried using custom MATLAB script. The deviation was calculated as sum of deviations of all involved area per lipids at given pressure value. The excess free energy of mixing was calculated by multiplying the mixing parameter by Avogadro number.

$$\Delta G = \int_{\pi_0}^{\pi} A_{12}d\pi - x_1\int_{\pi_0}^{\pi} A_1d\pi - x_2\int_{\pi_0}^{\pi} A_2d\pi \qquad (3)$$

$$\Delta G = \int_{\pi_0}^{\pi} A_{123}d\pi - x_1\int_{\pi_0}^{\pi} A_1d\pi - x_2\int_{\pi_0}^{\pi} A_2d\pi - x_3\int_{\pi_0}^{\pi} A_3d\pi \qquad (4)$$
